# Loss of Fmr1 reorganizes the multi-elemental composition across tissues in Fragile X Syndrome mice

**DOI:** 10.64898/2026.01.27.702117

**Authors:** Sabiha Alam, Jamie T. Reeves, Puni Jeyasingh, Shawn M. Wilder, Elizabeth A. McCullagh

## Abstract

Fragile X Syndrome (FXS) results from a genetic mutation which silences the expression of Fragile X Messenger Ribonucleoprotein (FMRP). FMRP serves various roles regulating cellular protein synthesis including mRNAs that code for proteins regulating ion flux. However, there are few studies measuring the elemental balance between FXS genotypes and tissues. Here, we measured the multivariate balance of 10 elements in tissues of wild-type and Fmr1-knockout mice to compare elemental composition of brain and somatic tissues within and across genotypes. Using a Bayesian mixed model approach, we found that the main differences between groups were between tissues, with significant effects of genotype and interaction of tissue on genotype. Wild-type feces were significantly higher in magnesium and sodium than knockout. Fur was significantly higher in potassium in wild-type, which was supported by the interaction effect of genotype with tissue. These results align with previous work showing FXS pathologies alter electrolytic and metal ion regulation, neuronal excitability, and gastrointestinal function. Future work should additionally test how elemental differences relate to function at the cellular level, as well as patterns of individual intake, digestion, assimilation, and/or excretion.

## Introduction

Fragile X Syndrome (FXS) is a neurodevelopmental disorder which results from a mutation in a single gene, *Fmr1* (Fragile X Messenger Ribonucleoprotein 1), on the X chromosome [1,2]. Mutation of this gene reduces the expression of its protein product, Fragile X Messenger Ribonucleoprotein (FMRP) [3,4]. Because FMRP is an RNA-binding protein, it affects a wide range of biological processes, including gastrointestinal function, synaptic activity, and neural development, among others. The diversity of these effects makes it difficult to catalog all downstream consequences of *Fmr1* disruption at the molecular level. Because all biological systems (e.g., cell, tissue) are made up of approximately 20 chemical elements (depending on structure, and process; [5], the relative abundance of all elements encompassing a system (i.e., the ionome) are appropriately conceptualized as a working unit [6,7]. Responses of the ionome to genetic perturbations have been used to advance understanding of various phenotypes [8] including disease [9–13] which could be considered complementary to other techniques such as transcriptomics [14,15] and proteomic analyses [16,17].

*Fmr1* encodes an RNA-binding protein (FMRP), and its loss affects diverse aspects of cellular physiology by disrupting post-transcriptional regulation of many target mRNAs. FMRP associates with ribosomes and polyribosomes to control the translation of transcripts involved in synaptic signaling, ion channel regulation, and metabolic processes [18,19]. Consequently, Fmr1 disruption alters both the synthesis and localization of proteins responsible for ion transport and storage (e.g., voltage-gated channels, metalloregulatory enzymes; [20–22]. Thus, FXS may secondarily influence elemental balance across tissues. These alterations in translational efficiency and post translational modifications can cascade into systemic changes in ionomic composition, potentially influencing neuronal function.

Individuals with FXS are often diagnosed with autism spectrum disorder (ASD), and exhibit overlapping clinical features such as antisocial behavior, communication deficits, cognitive impairments, and language delays [23]. Several studies have identified associations between ASD and specific elements: supplementation with zinc promotes restoration of synaptic proteins such as Shank3 and Shank2 and helps restore excitatory–inhibitory balance [24]. Magnesium, particularly in combination with vitamin B6, further mitigates neurobehavioral disorders in ASD [25,26]. Moreover, recent research demonstrates that ASD-related symptoms extend beyond neural dysfunction, including disruptions in balance across gut and peripheral tissues (iron -[27,28]; zinc - [29]; various trace metals - [30]. However, focusing on single elements or tissues, while informative, risks missing the broader systemic interactions that emerge from the coordination of multiple elements. Without a comprehensive ionomic perspective, the integrative patterns linking neural, gastrointestinal, and metabolic functions may remain unresolved.

In this context, we conducted an experiment to examine how the multielemental composition, the ionome, of neural and somatic tissues as well as digestive biproducts (feces) differs in the context of FXS between wild-type and knockout strains of mice. Because elemental concentrations are interdependent, these data were analyzed within a compositional framework that captures relative, rather than absolute, changes among elements. Using this approach, we explored the potential influence of *Fmr1* loss on overall elemental balance among brain regions and somatic tissues. We hypothesized that male mice lacking *Fmr1* would exhibit distinct multielement compositions relative to wild-type males. To test this hypothesis, we quantified and compared the ionomic composition of gut, brain, and noninvasive tissues such as fur and feces between genotypes. This compositional data analysis provides insight into systemic elemental reorganization in FXS and identifies tissues that may serve as noninvasive proxies for elemental diagnostics in clinical contexts [31].

## Methodology

### Experimental Animals and Design

In our study, we used an Fmr1 KO (stock #003025, Fmr1 KO) mouse model which recapitulates some core symptoms of FXS patients along with C57BL/6J (stock #000664, B6) wild-type background (control animals), which were obtained from the Jackson Laboratory and bred at Oklahoma State University [32]. All animals included in this study were male WT (n = 8) and KO (n = 6) mice ranging from 99–156 days of age at the time of tissue collection. Experimental groups were age-matched as closely as possible to minimize potential age-related effects on tissue elemental composition. Animals were group housed with littermates and generated through genotype exclusive mating (wildtype to wildtype and FXS to FXS). The results therefore reflect data from two cages of animals per genotype. All mice were on a 12-hour light cycle (6 AM- 6 PM) and received the same chow (LabDiet 500, Lab Supply, Fort Worth, TX) and bedding (alpha-dri, Innovive, San Diego, CA) and additional nestlets (cotton NST, Lab Supply, Fort Worth, TX). Since FXS is an X-linked trait and therefore more common in males, we limited the study to male animals. All experimental procedures were conducted under appropriate laws and NIH guidelines and our study principles received approval from the Oklahoma State University Institutional Animal Care and Use Committee (protocol 20-07).

### Tissue Collection and Preparation

Feces were collected from animals as they were passively in the holding cage, or we gently held the mice by the base of their tails and waited around 5 minutes until they defecated. After defecation, we used forceps to collect fresh fecal pellets and immediately placed those pellets into pre-labeled sterile tubes. Next, we euthanized mice by exposing them to isoflurane overdose. After confirming the death of the mice by lack of respiration and toe pinch reflex, we decapitated them and harvested their brains. We placed brains in a Petri dish and dissected the brains into specific brain regions including olfactory bulb and striatum. After removing the above-mentioned dissected brain regions, we pooled tissue from the pons, medulla, hippocampus, thalamus, hypothalamus as brain PMHTH (Figure S1). Since the brains were dissected fresh, these segregations (or pooling) indicated regions we were confident were isolated from other regions. We then transferred all dissected brain regions into a 2 mL Eppendorf tube for subsequent processing. Additionally, we collected samples from cecum and included fur for each individual mouse. To collect the cecum, we opened the abdominal cavity with a midline incision from the lower abdomen to the sternum, which exteriorized the intestine to expose the cecum—a large, pouch-like structure located at the junction between the ileum and the colon. We immediately transferred the cecum to a sterile, 2 mL Eppendorf tube. Later, to separate the cecal contents from the cecum tissue, we opened the cecum longitudinally on a sterile Pedri dish, then collected cecal contents and discarded the cecal tissue. For fur samples, we used clean, sterile scissors to trim the fur from the abdomen. Then, we transferred the clipped fur into a pre-weighed and labeled Eppendorf tube using sterile tweezers.

### Ionome Measurement

Once tissues were collected and stored, we dried them at room temperature in an oven at 55 °C for 72 hours. Then, we homogenized dried tissues using a mortar and pestle and weighed 4 – 12 mg for each sample. We digested homogenized samples using 100% trace metal grade HNO3 and 100% trace metal grade H2O2 in a 2:1 ratio; allowing them at least 24 hours for complete digestion before performing elemental analysis using inductively coupled plasma optical emission spectroscopy (ICP-OES, iCAP7400; ThermoScientific, Waltham, MA). We measured the concentrations of 10 biologically relevant elements in brain tissue using ICP-OES: Calcium (Ca), Copper (Cu), Iron (Fe), Potassium (K), Magnesium (Mg), Manganese (Mn), sodium (Na), Phosphorus (P), Sulfur (S), and Zinc (Zn). We retained wavelengths with 90% of measured samples within the limits of detection established by standard curves and replaced values over and under limits of detection with upper and lower limits. Then, we averaged emittance of multiple wavelengths when more than one wavelength quantified an individual element.

Of the 84 total samples analyzed for 10 elements (840 readings), 4 readings of sulfur, and 8 readings of zinc were missing values. Missing sulfur values included WT olfactory bulb #170, KO olfactory bulb #157, WT fur #162, KO PMHTH #142. Missing Zn values included KO PMHTH # 142 and 157, WT fur # 150, KO fur #151, WT olfactory bulb #169 and 170, KO olfactory bulb # 157 and 158. Thus, we retained *n*_WT_ = 6 and *n*_KO_ = 4 for sulfur, and *n* = 3 for zinc in both genotypes.

To account for missing values, we imputed missing values “impCoda” function from the “robCompositions” R package, which is specifically designed for compositional data [33]. A few missing values are common in ICP-based elemental analyses due to matrix effects and instrument detection limits [34]. Tissue elemental concentrations represent compositional data and were transformed to account for their inherent constraints. Concentrations of each element (µg/mg) were converted into percentages. We calculated the fill value (Fv) as the remaining unmeasured percentage of samples (i.e., 100 – sum of all measured elements). For each element, we then calculated its proportion of the total remaining unmeasured mass (Element/Fv) to standardize comparisons across samples in log space (i.e., additive log ratios; ALRs). These proportional values (Element/Fv) represent the relative abundances of each element within the measured elemental pools that are suitable for robust statistical analysis and inference (Greenacre, 2021).

### Mixed Modeling

We used a Bayesian multivariate mixed linear model to evaluate the effects of genotype (WT vs KO) and tissue (i.e., feces, striatum, cecum, olfactory bulb, fur, and PMHTH) on patterns of observed elemental variance. We used R package ‘brms’ to conduct the mixed modeling workflow (Bürkner, 2017) and used default values for all prior distributions. We conducted all analyses using statistical program R ver 4.5.2 (R Core Team, 2025).

We modeled each observation (*y_i_*) as a multivariate normal response variable with identity link, linear predictor *f*(η*_i_*) and deviance (σ) (Equation 1). An observation represents one measurement of 10 elements (*y_i_* = [*y_i_*Ca, *y_i_*Cu, *y_i_*Fe, … *y_i_*Zn]) from a single tissue within an individual, with expectation (η*_i_*) and residual deviance (σ). Although we sampled six tissue types, we chose not to include a flexible deviance parameter for each tissue due to limitations in sample size (*n_WT_* = 8; *n_KO_* = 6).

The linear predictor (*η_i_*) includes the design vector (*X_i_*) which maps population-level coefficients (tissue, genotype, and tissue x genotype interaction) onto individual observations (Equation 2). The linear predictor also includes the vector of element-wise group-level coefficients (*μ_j[i]_*) for observation *i* of individual *j* (i.e., random effect of individual on baseline concentrations across all tissues; Equation 3). Group-level coefficients were derived from a multivariate normal distribution centered at zero with covariance matrix (Σ*_μ_*), which is computed as the squared deviance of random element intercepts (σ*_μ_^2^*) times the correlation matrix of all elements (R*_μ_*; Equation 4).

The residual error term (□*_i_*) is a multivariate normal variable centered at zero with covariance matrix (Σ_□_ - Equation 5). Residual error includes all variance not explained by the linear predictor. The residual covariance matrix, like that in the group-level coefficients, was computed as the product of the squared residual deviance of random element intercepts (D_□_*^2^*) times the residual correlation matrix of all elements (R_□_ - Equation 6).

As is standard in the ‘brms’ package, we initialized a Markov Chain Monte Carlo (MCMC) sampler with 4 chains of 2,000 iterations each. In each chain, we used the standard burn-in of 1,000 iterations and retained 1,000 iterations from each chain. Thus, the maximum possible effective sample size was 4,000 for each parameter.

### Model Validation

To evaluate MCMC convergence, we assessed whether rank-normalized split Gelman-Rubin diagnostics (□) for all estimated parameters met the validation threshold of □ < 1.05 (Vehtari et al., 2021). Additionally, we evaluated the effective sample sizes of model parameters, including the bulk and tail measures (*ESS*_bulk_ and *ESS*_tail_). These measures quantify the exploration of high-probability and tail posterior parameter spaces by all chains. We used a standard threshold of *ESS*_tail_ > 400 and *ESS*_bulk_ > 400 to assess chain convergence and stability.

After obtaining model results, we performed posterior predictive checks and assessed residual dispersion. Posterior predictive checks allow visual comparison of the observed and predicted distributions. We performed posterior checks only for elements which displayed genotype-level contrasts in at least one tissue with 95% Highest Posterior Density (HPD) interval not overlapping zero. To assess residual dispersion, we extracted residuals of conditional predictions for all samples and stratified residual error post-hoc based on observation-level genotype and tissue factors. Then, we calculated the standard deviation (SD) and interquartile range (IQR) of residuals for each group and compared values across groups.

### Model Evaluation

We computed the posterior R^2^ distributions for each element to quantify explanatory power and related uncertainty. Then, we inverted the residual covariance matrix (Σ_□_) to obtain the precision matrix (Ω_□_ = Σ_□_ ^-1^) and extracted isolated element-wise error as the inverse of values on the diagonal of the precision matrix (i.e., element error independent of covariance). We thus decomposed posterior variance into population-level components (tissue, genotype, and tissue x genotype interaction), group-level components (individual), shared residual error (elemental covariance), and isolated residual error.

We performed post-hoc contrasts of predicted values across experimental groupings to compute estimated marginal means. We performed pairwise contrasts across all population-level coefficients (tissue, genotype, tissue x genotype) by directly computing the difference between predicted group-level values across each iteration in retained samples of all four chains. We consider contrasts indicative of an observed effect when 95% HPD intervals do not cross zero and report the posterior mean magnitude and direction of the effect on the log-ratio scale.

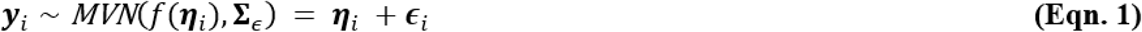

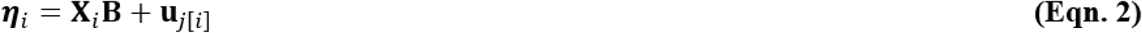

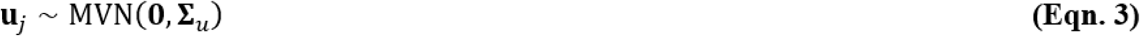

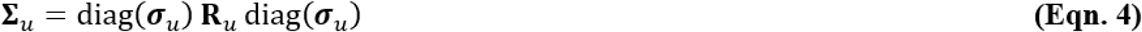

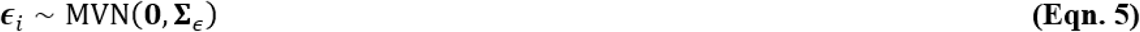

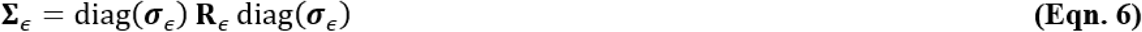

## Results

### Global model

Overall, our model demonstrated strong evidence of convergence (maximum□= 1.006) and stability (minimum *ESS*_bulk_ = 514, *ESS*_tail_ = 880; Table S1). In addition, predictive distributions were generally well-aligned with observations (Figure S1, S2, Table S2). Residual dispersion from posterior predictions (SD and IQR) did not differ between KO and WT mice (Table S3), but Striatum demonstrated significant overdispersion relative to other tissues (∼ 4 times higher; Table S4). Estimates and credible intervals for model parameters are available in Table S5 and S6).

The overall model predicted effects of genotype, tissue, and interaction on elemental ratios (Figure 1a, 1b). Total variance explained by population-level (fixed) effects varied among elements, but median estimates exceeded 0.75 in 70% of elements (Table 1). The population-level effects explained nearly all variance in Mg and Mn (median 98 and 99%, respectively) with high confidence. However, Ca, S, and Zn generally were not reliably predictable, as median estimates were between 0.28 and 0.46 and uncertainty was substantially higher than other elements (Table 1).

**Figure 1a.**
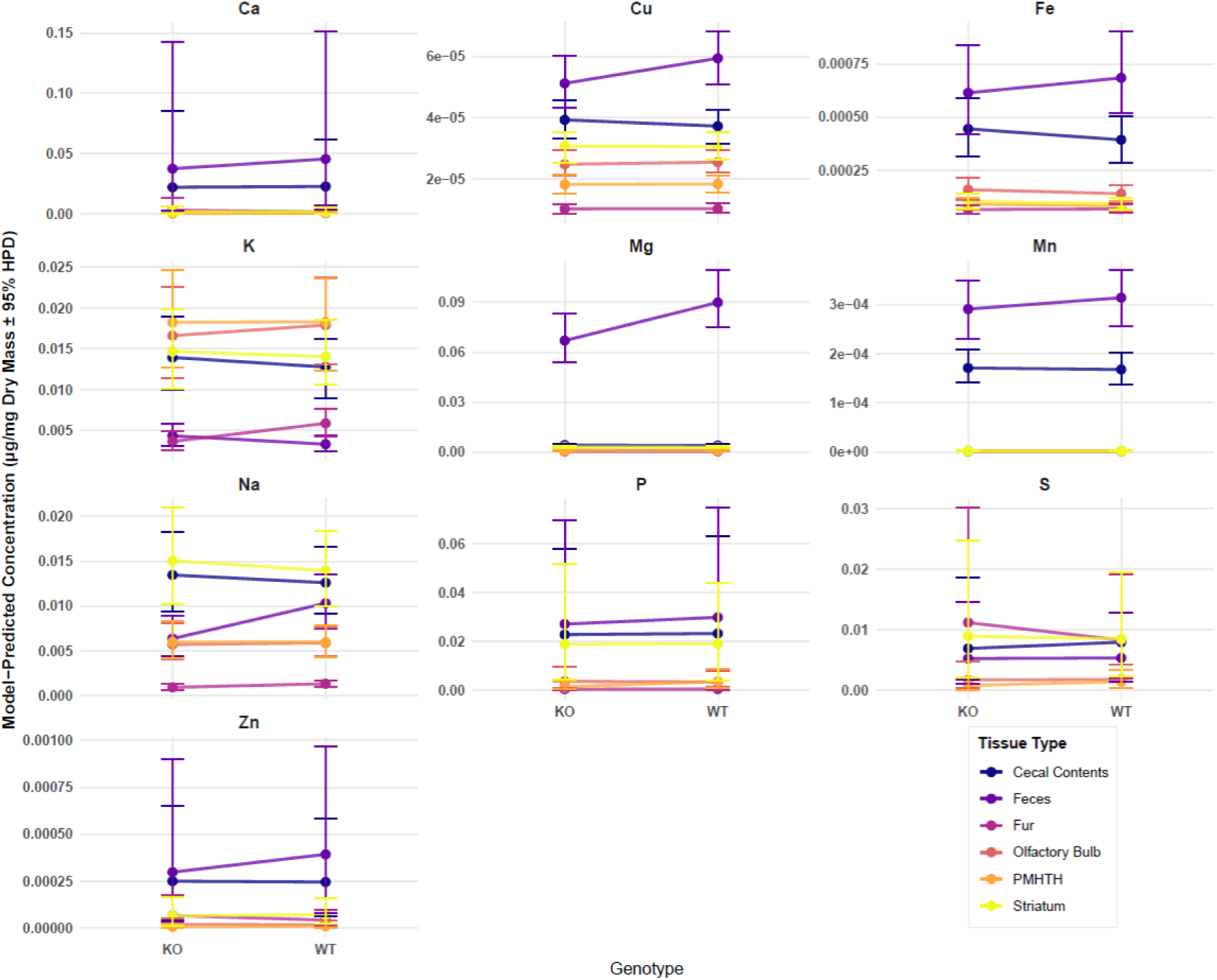
Estimated marginal concentration profiles across tissues and genotypes. Facets track model-predicted elemental concentrations (µg/mg dry mass) back-transformed from the multivariate log-ratio scale via exponentiation. Points denote posterior medians, and error bars represent the 95% highest posterior density (HPD) intervals.

**Figure 1b.**
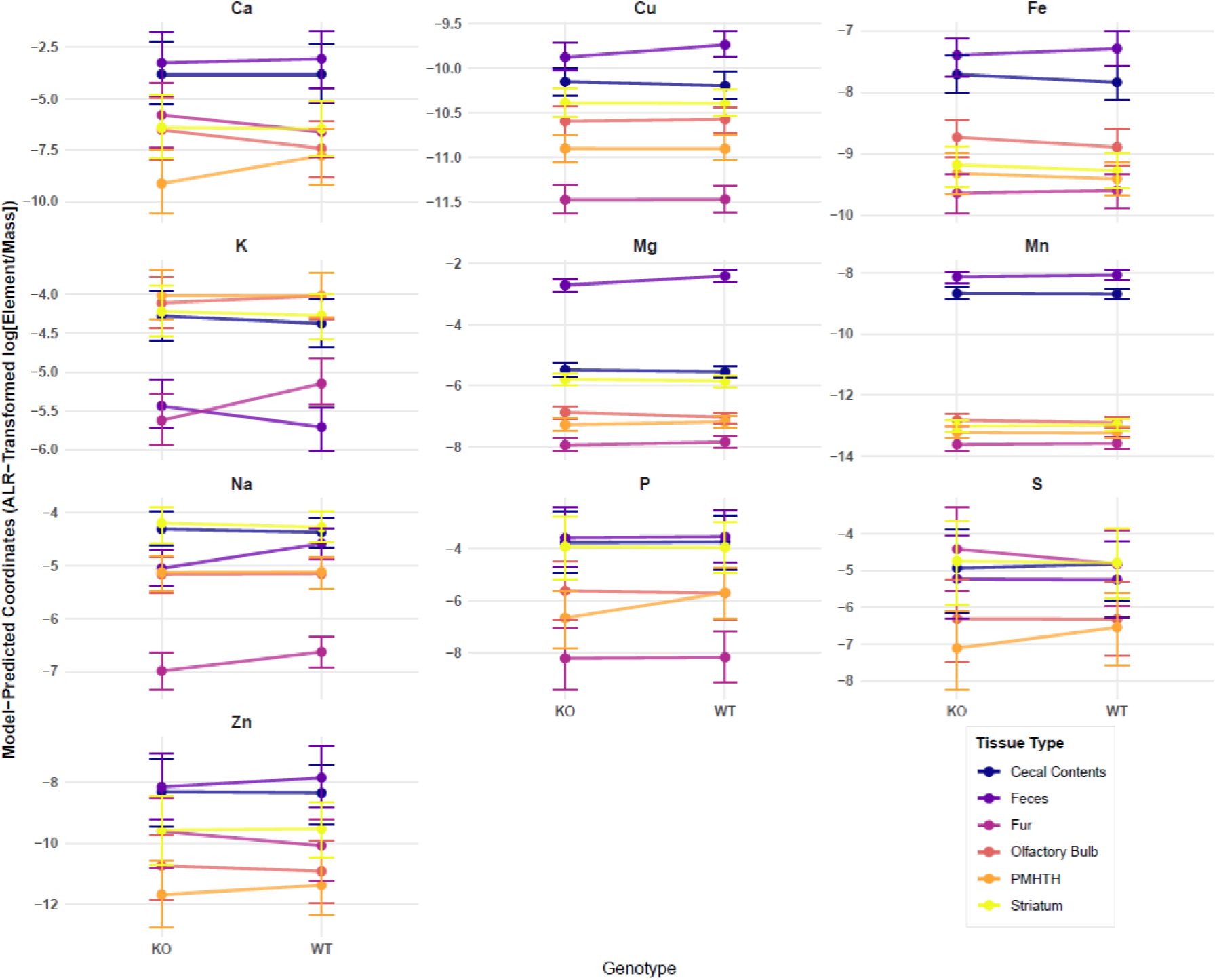
Estimated marginal concentration profiles across tissues and genotypes. Facets track model-predicted elemental ratios (ALR-transformed). Points denote posterior medians, and error bars represent the 95% HPD intervals.

**Table 1.** Predicted explanatory power (R-squared distribution) of total population-level (i.e., fixed) effects on individual elements. R-squared was iteratively calculated from posterior predictions using 4,000 draws for each element. Columns represent median estimate, estimated error, and quantiles.

| Element | Estimate | Est.Error | Q2.5 | Q97.5 |
| --- | --- | --- | --- | --- |
| Ca | 0.459 | 0.051 | 0.348 | 0.544 |
| Cu | 0.922 | 0.008 | 0.905 | 0.935 |
| Fe | 0.845 | 0.015 | 0.809 | 0.869 |
| K | 0.774 | 0.025 | 0.716 | 0.814 |
| Mg | 0.983 | 0.001 | 0.979 | 0.985 |
| Mn | 0.992 | 0.001 | 0.990 | 0.993 |
| Na | 0.820 | 0.018 | 0.780 | 0.847 |
| P | 0.557 | 0.043 | 0.458 | 0.628 |
| S | 0.281 | 0.057 | 0.163 | 0.384 |
| Zn | 0.436 | 0.052 | 0.323 | 0.529 |

Tissue in general explained the most variance in elemental ratios (especially in Mg and Mn), but genotype and tissue-genotype interaction also influenced some elements (Figure 2). Group-level (i.e., random) individual mouse-level coefficients explained very little observed variance in any element, but some elements displayed substantial isolated and correlated residual error (Figure 2; Table 2a, b).

**Figure 2.**
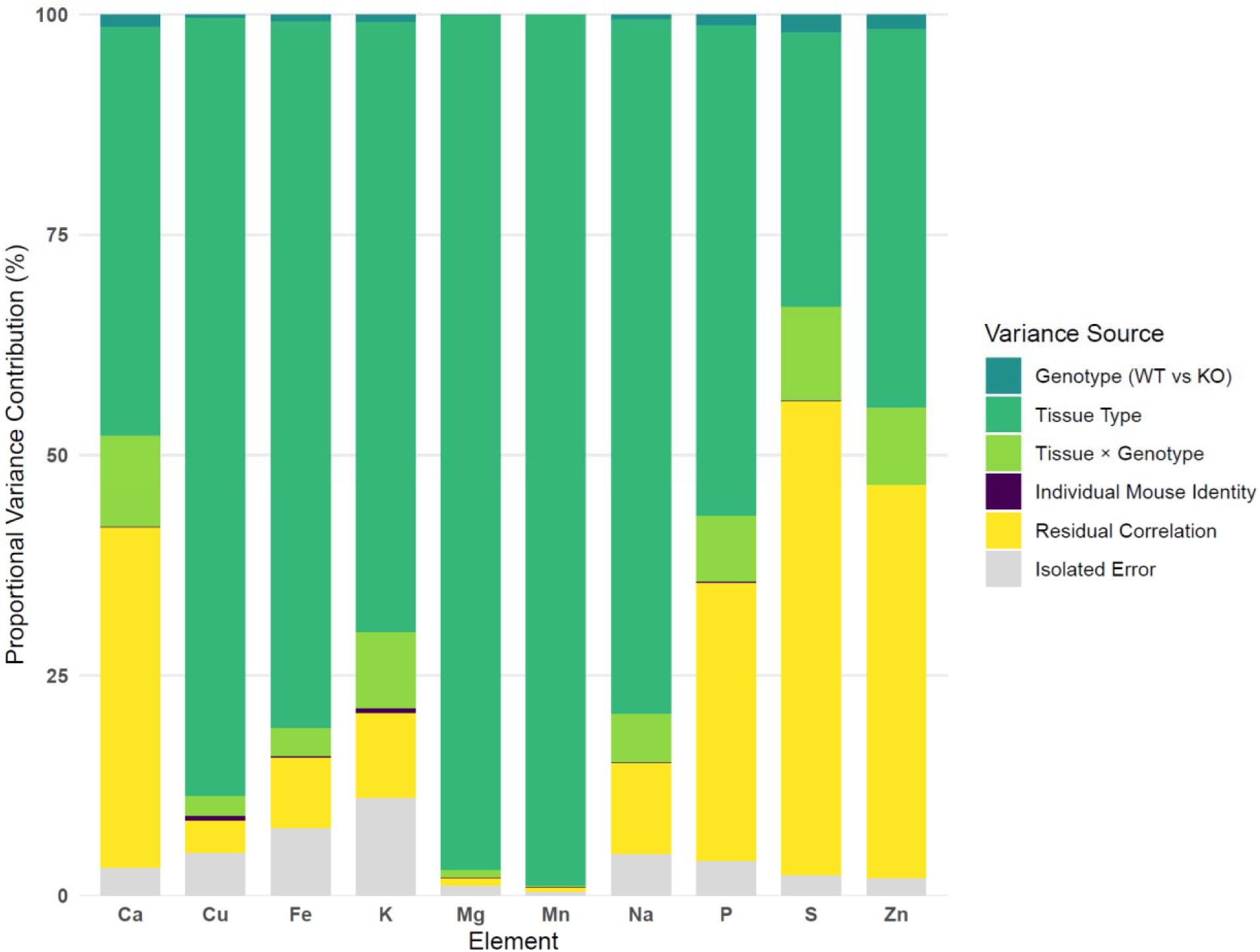
Global variance decomposition across elements. Stacked bars illustrate the relative percentage of total variance allocated to distinct hierarchical model components for each element. To account for the mathematical closure properties of compositional data, residual variance is partitioned into isolated error (grey) and residual correlations (yellow) using a Squared Multiple Correlation (SMC) projection of the residual matrix. All profiles were normalized to a strict 100% variance closure pool per element.

**Table 2.** A) Global variance decomposition showing percentage contributions of experimental factors to total ionomic variance observed in original data. Fixed effects include Genotype, Tissue, and Genotype x Tissue Interaction (Interaction). Random effects include mouse identity (Individual). Residual variance is partitioned into isolated unexplained error and shared pairwise variance allocations calculated using Squared Multiple Correlation (SMC). B) Percentage contribution of residual correlations between individual pairs of elements.

**A)**
| Element | Genotype | Tissue | Interaction | Individual | Isolated Error | Residual Correlation |
| --- | --- | --- | --- | --- | --- | --- |
| Ca | 1.4 | 45.5 | 10.2 | 0.1 | 3.1 | 37.9 |
| Cu | 0.4 | 87.6 | 2.2 | 0.6 | 4.8 | 3.6 |
| Fe | 0.8 | 79.2 | 3.1 | 0.2 | 7.5 | 7.9 |
| K | 0.9 | 68.0 | 8.5 | 0.5 | 10.9 | 9.5 |
| Mg | 0.1 | 96.9 | 0.8 | 0.0 | 1.1 | 0.9 |
| Mn | 0.0 | 98.8 | 0.2 | 0.0 | 0.4 | 0.4 |
| Na | 0.6 | 78.0 | 5.5 | 0.1 | 4.6 | 10.2 |
| P | 1.2 | 54.6 | 7.2 | 0.2 | 3.8 | 30.9 |
| S | 2.0 | 30.4 | 10.4 | 0.1 | 2.2 | 52.4 |
| Zn | 1.7 | 42.0 | 8.6 | 0.1 | 2.0 | 43.6 |

**B)**
| Element | Ca | Cu | Fe | K | Mg | Mn | Na | P | S | Zn |
| --- | --- | --- | --- | --- | --- | --- | --- | --- | --- | --- |
| Ca | 0.0 | 0.4 | 0.4 | 0.3 | 8.8 | 0.6 | 1.5 | 17.1 | 5.5 | 0.5 |
| Cu | 0.1 | 0.0 | 0.1 | 0.2 | 0.0 | 2.3 | 0.2 | 0.1 | 0.0 | 0.1 |
| Fe | 0.2 | 0.2 | 0.0 | 0.2 | 0.2 | 0.8 | 4.4 | 0.2 | 0.1 | 0.2 |
| K | 0.2 | 0.4 | 0.3 | 0.0 | 0.5 | 0.6 | 4.7 | 0.2 | 0.5 | 0.6 |
| Mg | 0.3 | 0.0 | 0.0 | 0.0 | 0.0 | 0.0 | 0.2 | 0.1 | 0.1 | 0.0 |
| Mn | 0.0 | 0.2 | 0.0 | 0.0 | 0.0 | 0.0 | 0.0 | 0.0 | 0.0 | 0.0 |
| Na | 0.4 | 0.4 | 2.7 | 2.8 | 1.8 | 0.3 | 0.0 | 0.5 | 0.2 | 0.2 |
| P | 17.0 | 0.3 | 0.4 | 0.4 | 3.6 | 0.3 | 1.7 | 0.0 | 1.7 | 3.1 |
| S | 6.8 | 0.3 | 0.4 | 1.2 | 2.4 | 1.4 | 0.8 | 2.1 | 0.0 | 33.1 |
| Zn | 0.5 | 0.4 | 0.5 | 1.4 | 0.5 | 0.5 | 0.8 | 3.7 | 31.7 | 0.0 |

### Contrasts

Contrasts were performed to detect posterior differences between population-level (i.e., fixed) groupings found genotype-level effects within individual tissues for three elements: K, Mg, and Na (Figure 3). The posterior log-ratios of Mg and Na in feces were respectively 0.5 and 0.3 units lower in KO than WT mice (Figure 3). The posterior log-ratio of K in fur was 0.5 units lower in KO than WT mice (Figure 3).

**Figure 3.**
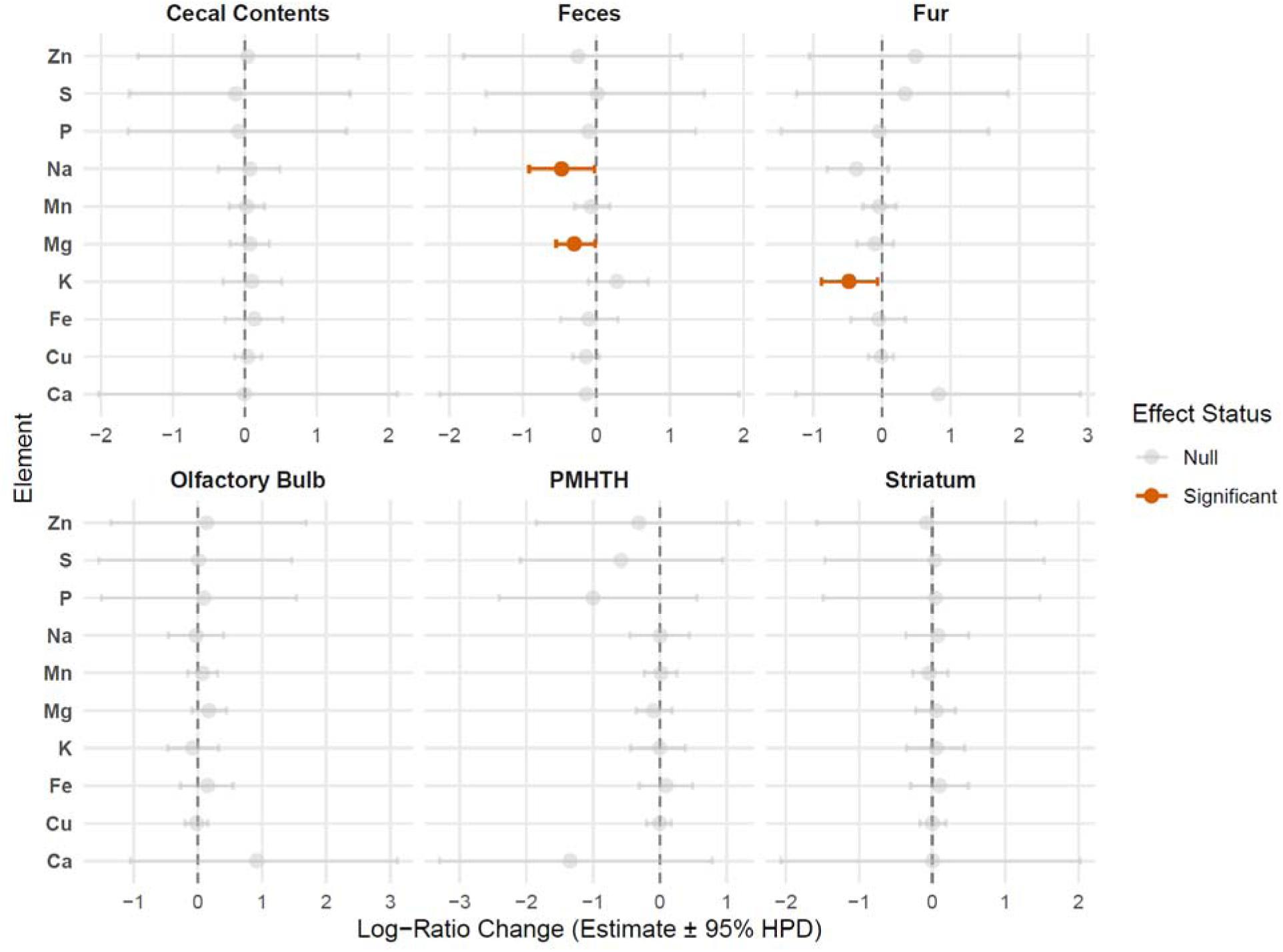
Pairwise contrasts of genotype effects on tissue-level ionomes. Points represent the median of the posterior difference distribution on the native additive log-ratio (ALR) scale. Horizontal bars represent the 95% HPD intervals for the pairwise differences between wild-type (WT) and knockout (KO) genotypes across tissue types.

All elements demonstrated at least one significant tissue-level contrast, but a substantial number of elements demonstrated multiple differences in log-ratio at the tissue level (Figure 1a, 1b, 4). However, only K displayed significant effects of the interaction between tissue and genotype (Figure 1a, 1b, 5). Further, pairwise contrast within individual mice of elemental intercepts failed to detect significant within-individual correlations between elements (Figure S3).

**Figure 4.**
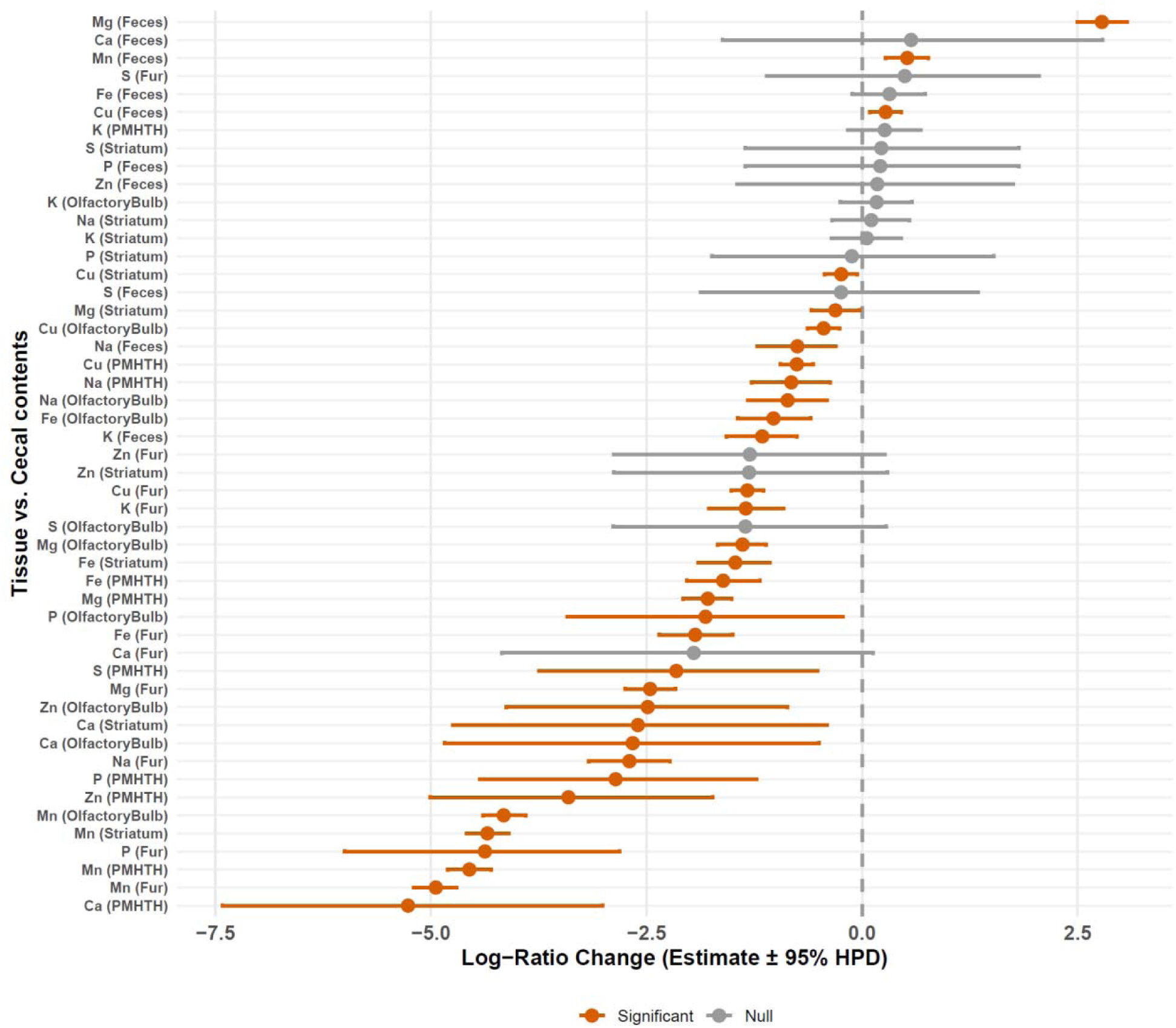
Global tissue fixed-effect coefficients on the log-ratio scale. Forest plot isolating the structural baseline shifts caused by anatomical tissue differentiation, independent of genotype. Coefficients represent the unique log-ratio shift relative to the designated reference tissue. Points denote posterior medians and horizontal bars indicate 95% HPD.

**Figure 5.**
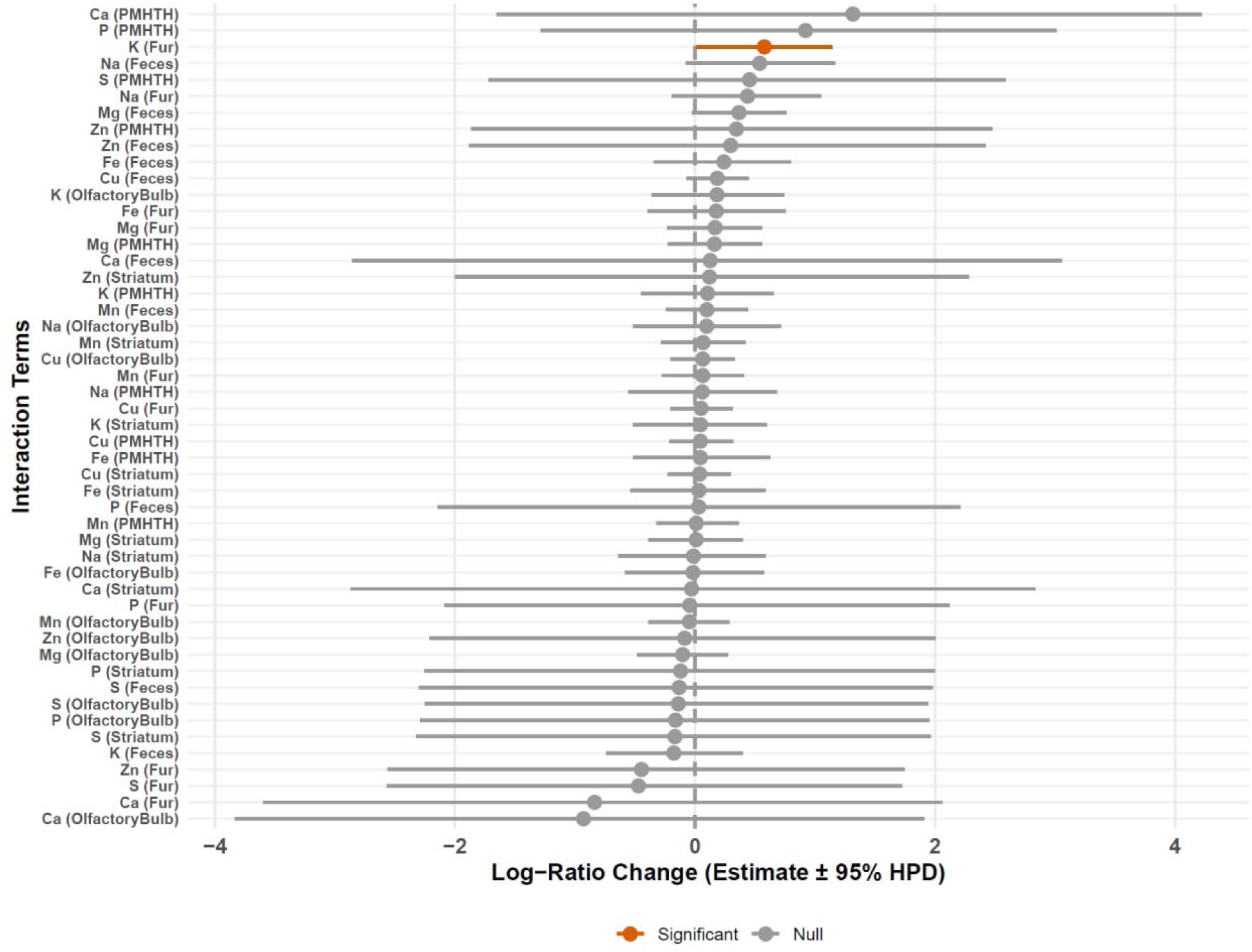
plot displaying the fixed-effect interaction terms (Tissue x Genotype) from the global joint model. Points denote posterior medians and horizontal bars represent the 95% HPD.

### Residuals

Predicted isolated error (from inverted covariance matrix) was generally low (< 10%) but reached a maximum in K and a minimum in Mn (Figure 2). However, residual correlation explained the most variance of any source in S and Zn (Figure 2). These two elements displayed the strongest residual correlation with each other, but other pairs of elements also displayed strong residual correlation, including Ca - P, Ca - S, Ca - Zn, P - S, and P - Zn (Figure 6).

**Figure 6.**
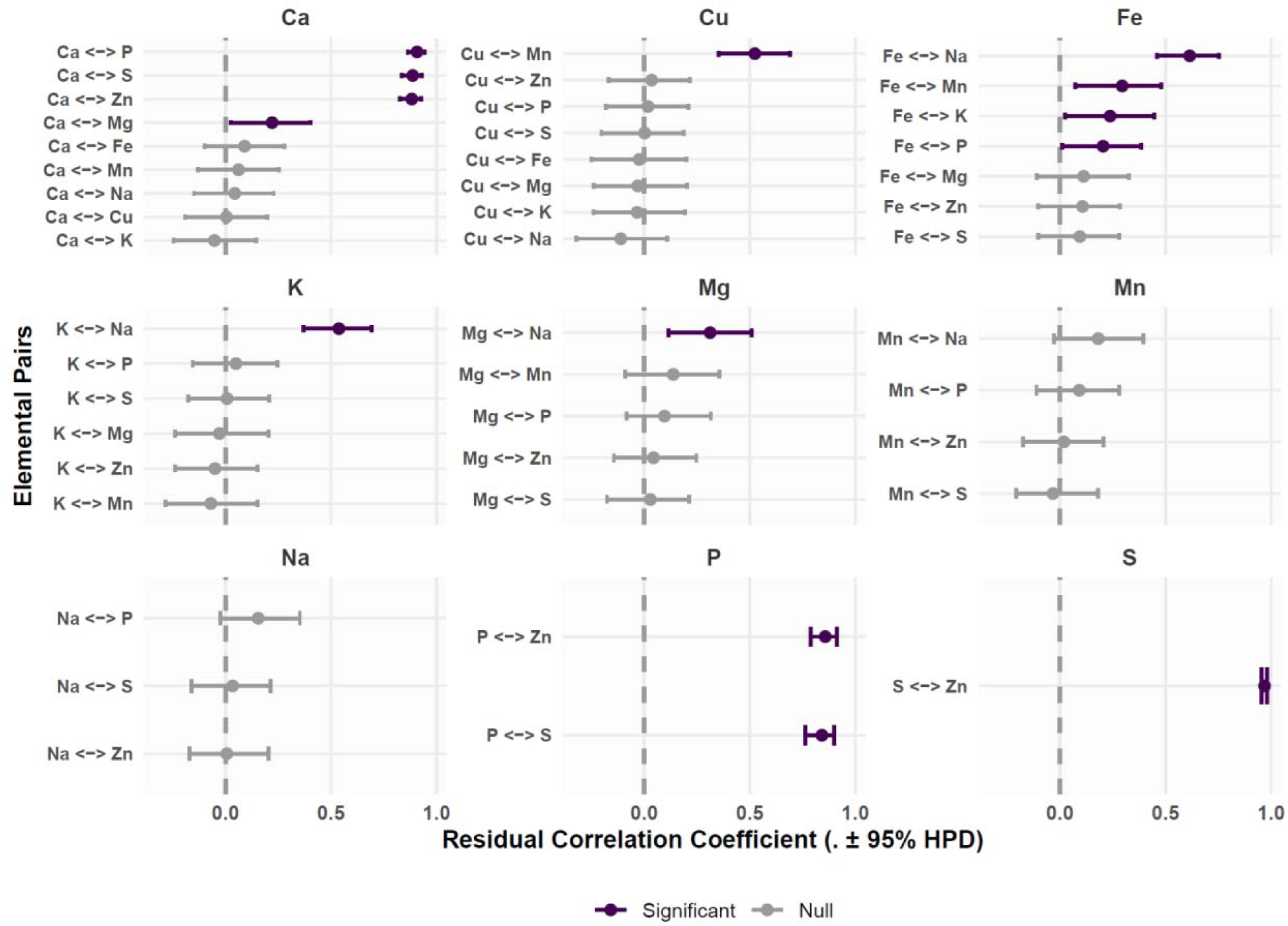
Faceted forest plots displaying the median coefficients and 95% HPD for residual correlation between pairs of elements. Values for each pairwise comparison are available in Table 1.

Post-hoc binning of posterior residual error by extracting correlation coefficients from MCMC iterations revealed differences in residual correlation among tissues (Figure S4). For example, residual correlation in Fe-P and Fe-K only occurred in fur. Alternatively, the residual Fe-Na correlation was higher and more certain in fur than that in olfactory bulb or PMHTH (Figure S4).

Post-hoc assessment of differences in elemental correlation also revealed differences among genotype (Figure S5). Correlations between Mg - Na, Fe - Mn, and Fe - K were moderately positive (median 0.3-0.4) for WT, but 95% HPD intervals of KO mice overlapped with 0 (Figure S5).

## Discussion

The results of this work provided limited support for our hypothesis that FXS affects the compositional ionome of ten elements (Ca, Cu, Fe, K, Mg, Mn, Na, S, P, and Zn) in neural and somatic tissues. In this study, multivariate analyses demonstrated that elemental composition patterns were driven mainly by tissue type rather than genotype, with by far the most variation attributable to tissue origin. However, multiple elements differed between WT and KO animals in feces (Mg, Na) and fur (K), as well as differences in residual correlation of elements among tissues and genotype. Notably, we found that the striatum had substantially higher residual dispersion than other tissues, indicating ∼4 times more elemental variance in this tissue. Further, high residual correlation and heavily weighted residual error in Ca, P, S, and Zn suggest that unaccounted factors drive a substantial proportion of variance in these elements, in some cases exceeding variance among tissues. Overall, these differences support previous observations that Fmr1-KO influences gut and broader epidermal function through electrolyte (e.g., Na, K, Mg) dysregulation [20,21,35,36].

We found several differences in elemental ratios between individual tissues of Fmr1-KO and WT mice. For example, we detected an interactive effect of tissue: genotype on potassium contribution, but mean potassium contribution differed between Fmr1-KO and WT only in fur. Interestingly, feces from WT mice contained relatively higher Mg and Na levels compared with Fmr1-KO mice, suggesting potential differences in elemental handling or excretion between genotypes. Because Mg and Na are important regulators of numerous cellular and neuronal processes, altered homeostasis of these elements may contribute to broader physiological dysfunction associated with Fmr1 deficiency.

Previous work further showed children with ASD self-select diets higher in Mg and lower in protein and Ca than children without ASD, and that selective diets were more likely to result in substantial deficiency of at least one nutrient [37]. Interestingly, evidence from children with ASD suggested Mg supplementation might alleviate neuro-behavioral symptoms [26]. Although this remains to be rigorously tested, previous authors advocate for the relevance of understanding Mg action in FMR1 pathways [38].

### Limitations and Future Directions

In conclusion, we found differences in ionomic composition between tissues in all 10 individual elements; although, we found genotype-level differences in overall composition and individual elements only in some tissues. Yet, several limitations must be acknowledged. First, our study did not use mechanistic assays to link elemental changes directly to FMRP function or downstream molecular pathways. Additionally, other elements contribute to tissue-level ionomes and are required for cellular processes and structures (e.g., carbon, nitrogen, silicon, etc.) which we did not measure here. As a result, the total percentage of original sample mass measured was in most cases quite small. Furthermore, although experimental groups were age-matched as closely as possible, age-related physiological and metabolic changes may also influence tissue elemental composition. Future studies with larger cohorts specifically designed to evaluate age-dependent variability in tissue ionomic profiles would therefore be valuable. Additionally we only used male mice in our study, previous work has shown that there are sex differences physiologically [39,40] and behaviorally [41] in FXS mice suggesting that sex may also be an important factor for future work. Although isoflurane exposure was applied uniformly across experimental groups, previous studies have reported that inhalation of anesthetics can affect iron homeostasis, ferroptosis-related pathways, intracellular metabolite diffusion, and mineral-regulatory processes [42,43]. Together, these findings suggest that isoflurane may affect elemental and metabolic homeostasis and therefore represents a potential confounding factor when interpreting tissue mineral composition data, particularly iron-related measurements and is an important difference between our work and previous published literature [28]. Additionally, in this study, all tissues were collected without any vascular perfusion, and serum elemental composition was not measured; therefore, tissue ionomic profiles may partially reflect circulation-associated elemental content in addition to tissue-specific organization.

While previous work has also measured elemental differences in FXS, direct quantitative comparison between the present study and others should be interpreted cautiously because the studies differed substantially in tissue selection, analytical methodology, and normalization strategy. Talvio et al. quantified absolute elemental concentrations (µg/g wet weight) in cortex, cerebellum, heart, liver, and spleen using ICP-MS, whereas our study analyzed dried tissue samples using ICP-OES, which is less sensitive, within a compositional framework focused on relative multielement organization across olfactory bulb, striatum, PMHTH, fur, feces, and cecal contents [28]. Additionally, the studies differed in tissue preparation and experimental workflow, including anesthetic exposure, tissue pooling strategy, drying procedures, digestion workflows, and elemental normalization approaches. These methodological differences are expected to influence absolute elemental concentrations, particularly for redox-sensitive elements such as iron. Importantly, Talvio et al. similarly emphasized cautious interpretation due to limited statistical power and multiple comparisons [28]. Our study was designed to evaluate whether loss of Fmr1 alters relative multielement organization across neural and somatic tissues. In this compositional framework, interpretation primarily depends on relative elemental relationships rather than direct equivalence of absolute concentrations.

Nevertheless, analyzing multi-tissue, multi-elemental data allowed us to identify systematic and genotype-dependent ionomic shifts in the Fmr1 knockout male mouse model. The combined use of additive log-ratio (ALR) transformation and compositional data analysis provided a robust framework for handling the constrained and interdependent nature of multi-elemental data [44]. Genome-wide RNAi ionomics screens in human cells have further demonstrated that multielemental profiles can reveal previously unrecognized genes and regulatory networks governing trace element metabolism [45], in the context of this study, ionomic profiling of non-invasive tissues such as feces or fur/hair could be linked to behavioral deficits enabling biomarker discovery.

Even though, in this study we measured overall relative concentration of an element in different neural as well as somatic tissues, future studies should assess not only total elemental abundance but also labile mineral content. The reason behind this logic is total tissue mineral levels do not unveil whether an element is biologically in active form, safely stored, protein-bound, redox-active, or localized within specific cellular compartments in tissues. This especially carries crucial insights for iron, because iron toxicity depends strongly on its biochemical form and intracellular localization. For example- ferritin-bound or transferrin-bound iron is relatively protected, whereas labile Fe² can promote Fenton chemistry, oxidative stress, lipid peroxidation, and ferroptosis [46]. Thus, two tissues may show similar total iron concentrations, yet differ substantially in toxicity or physiological impact if one contains a larger labile/redox-active iron pool. Therefore, measuring only total tissue iron could not fully capture biologically relevant alterations in iron homeostasis in our study.

For the other elements measured in this study — their labile pools may provide more mechanistic insight than total elemental concentration alone. For example, labile Ca²□ regulates ion transport, secretion, epithelial signaling, and cellular stress responses [47]; labile Zn²□ functions as a signaling ion [48]; and labile Cu and Mn may influence redox biology, mitochondrial function, and oxidative stress [49].

Future studies should investigate labile metal pools within the gut lumen and microbial environment, because total elemental abundance in luminal contents may not accurately reflect the fraction of minerals that is biologically available to microbes or host epithelial cells. Since the concept of “labile metals” refers to exchangeable, bioavailable metal ions that actively participate in microbial metabolism, redox reactions, signaling, and host–microbe interactions rather than metals that are tightly bound or inertly stored. Previous studies also emphasize that bacteria dynamically regulate labile metal pools to maintain metabolic homeostasis, oxidative stress resistance, and nutrient acquisition [50].

This is particularly relevant to our study because we quantified elemental concentrations in intestinal luminal contents, where metals are simultaneously influenced by diet, microbial utilization, epithelial transport, mucus interactions, and host inflammatory responses [51,52]. Therefore, future studies should examine not only total elemental levels but also the biologically active and compartment-specific metal pools, as these may better explain how altered mineral homeostasis contributes to gut microbiome changes and other neural as well as peripheral phenotypes in FXS. Future work is needed to combine bulk elemental profiling with approaches that measure bioavailable and compartment-specific mineral pools. This would help determine whether altered mineral composition in FXS reflects changes in total abundance, altered intracellular distribution, or shifts in the biologically reactive mineral pools most likely to affect mitochondrial metabolism, oxidative stress and cellular toxicity.

While challenging, future work integrating ionomic patterns with other biochemical systems (e.g., transcriptome, proteome) would help identify mechanistic pathways underlying loss of FMRP and disrupted elemental homeostasis. Validation of tissue-specific elemental signatures (e.g., Mg and Na in feces; K in fur) may further enable biomarker discovery and the development of ion-targeted therapeutic strategies for Fragile X Syndrome particularly if supplementation or severity could be linked with ionomic variation.

## Declaration of competing interests

The authors have no competing interests to declare.

## Notes

### Competing Interest Statement

The authors have declared no competing interest.

### Summary of Updates

We have updated the analysis and therefore most of the manuscript has changed.

### Funding

- National Institutes of Health: 5P20GM152333-02

