## supplemental info for "Loss of Fmr1 reorganizes the multi-elemental composition across tissues in Fragile X Syndrome mice"

**Supplementary Materials**

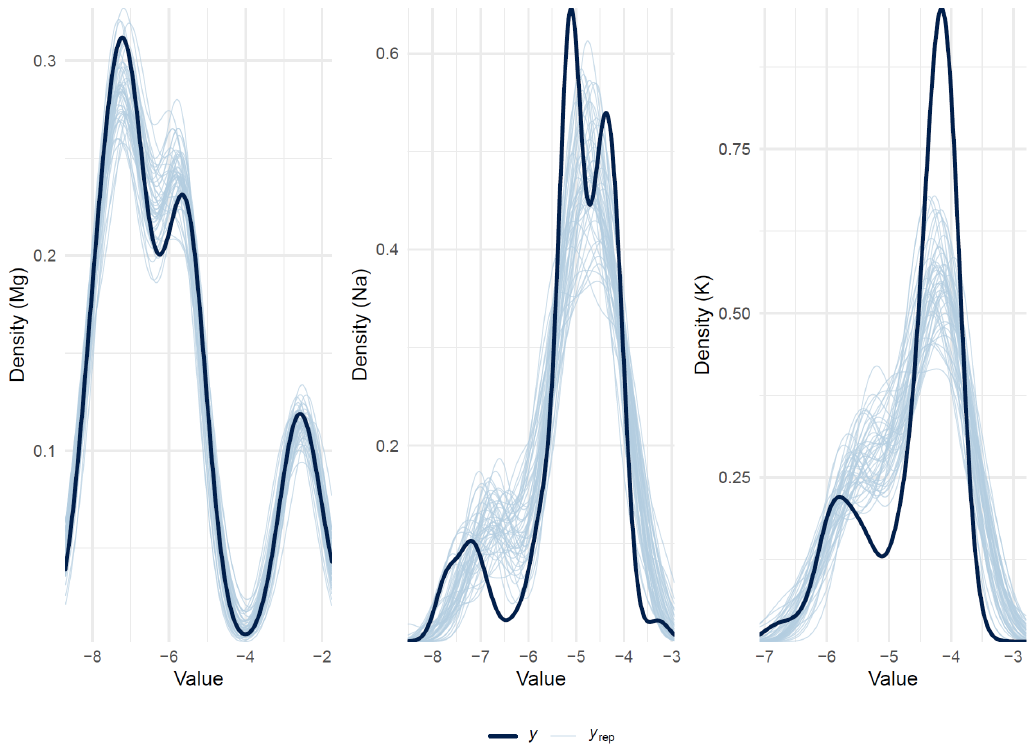

**Figure S1.** Multivariate posterior predictive checks (PPC) for core elemental indicators. Density overlays validating global model fit for representative elements (Mg, Na, and K). The solid black line *y* defines the empirical distribution of the observed log-ratio data; thin turquoise lines *y*_rep_ represent 50 independent simulated datasets generated from the posterior predictive distribution. High alignment between empirical and simulated densities verifies structural model adequacy and distributional assumptions.

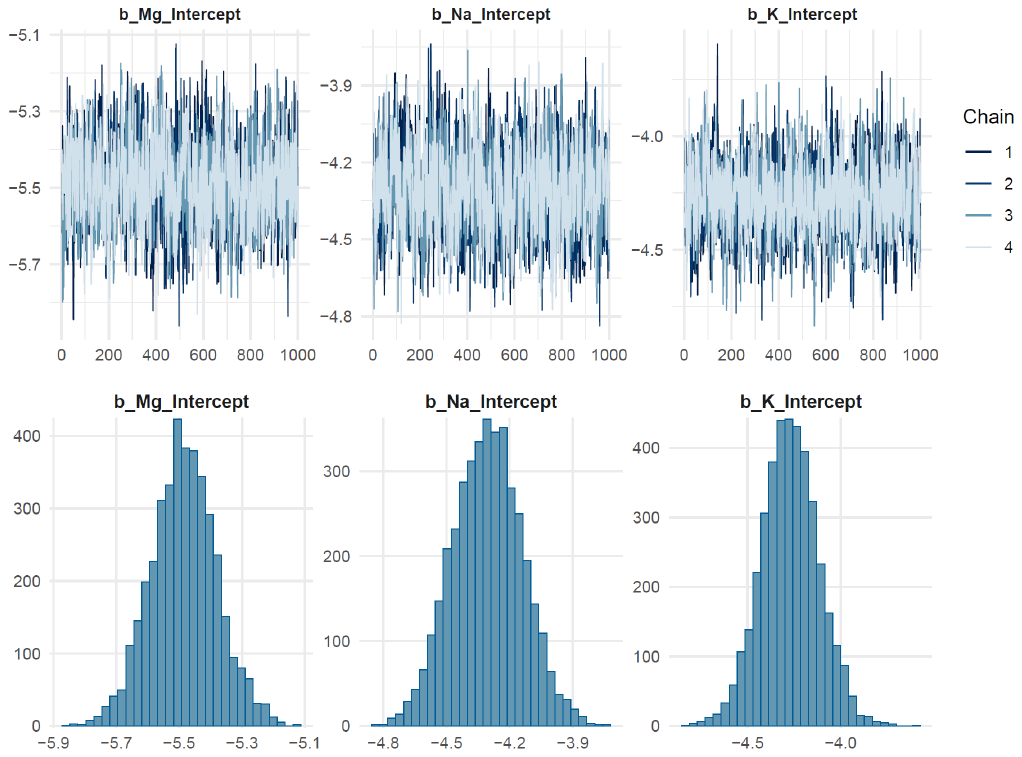

**Figure S2.** Markov Chain Monte Carlo (MCMC) sampling diagnostics. Standalone posterior histograms split by individual sampling chains for intercept parameters (Mg, Na, and K). Overlapping unimodal distributions across all four computational chains visually confirm robust parameter space exploration, chain mixing, and global convergence.

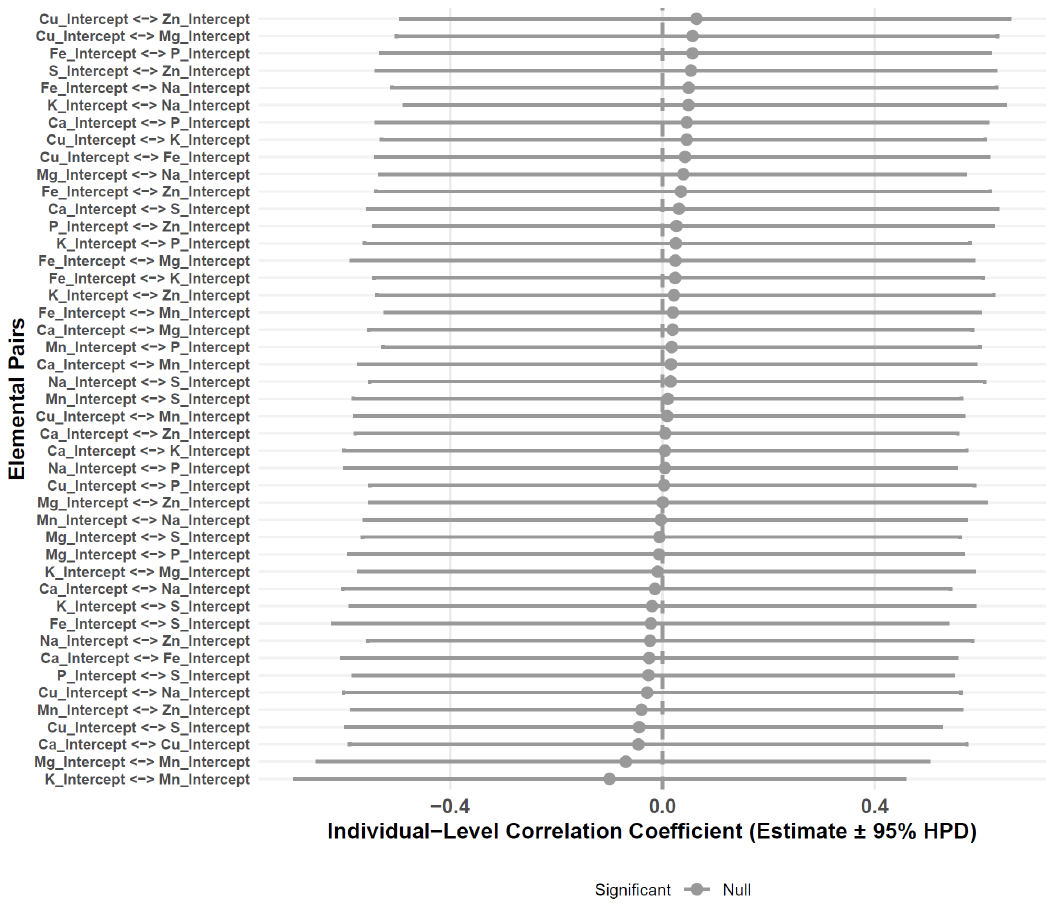

**Figure S3.** Forest plot displaying the median residual correlation coefficients and corresponding 95% HPD intervals for baseline elemental co-accumulation patterns among individual mice. Points represent median and lines represent 95% HPD.

**Figure S4 (Below).** Forest plot displaying the tissue-specific median residual correlation coefficients and corresponding 95% HPD intervals. Points represent median and lines represent 95% HPD.

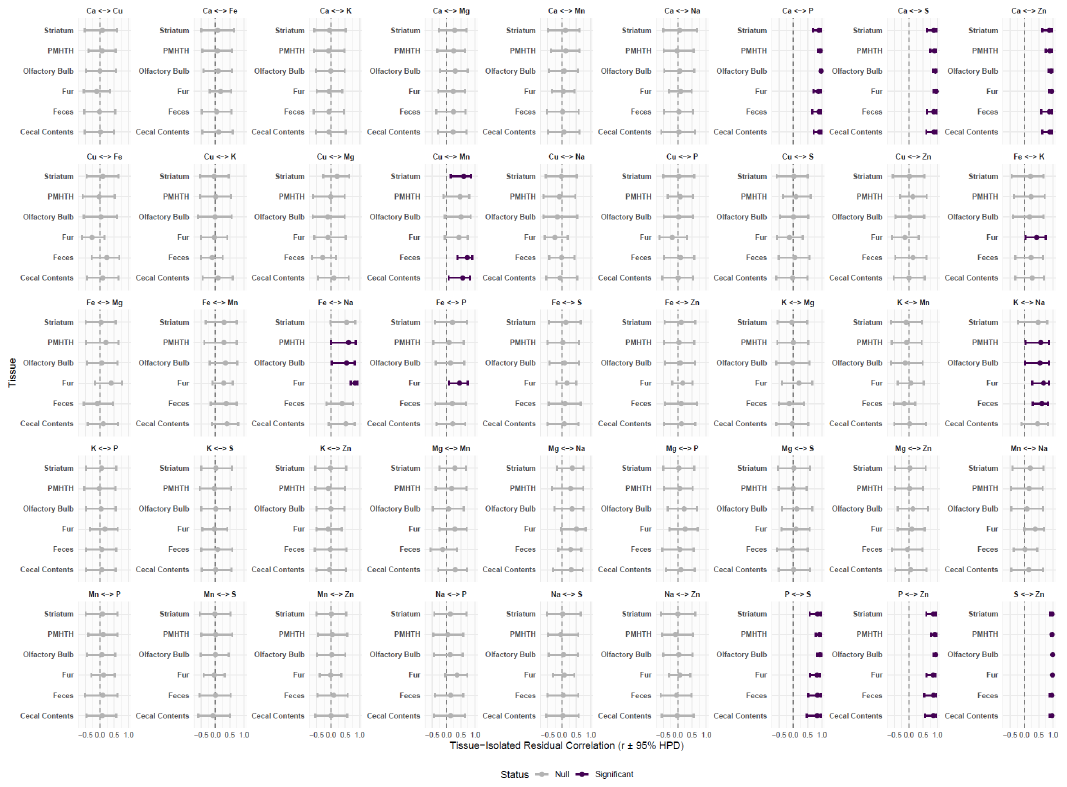

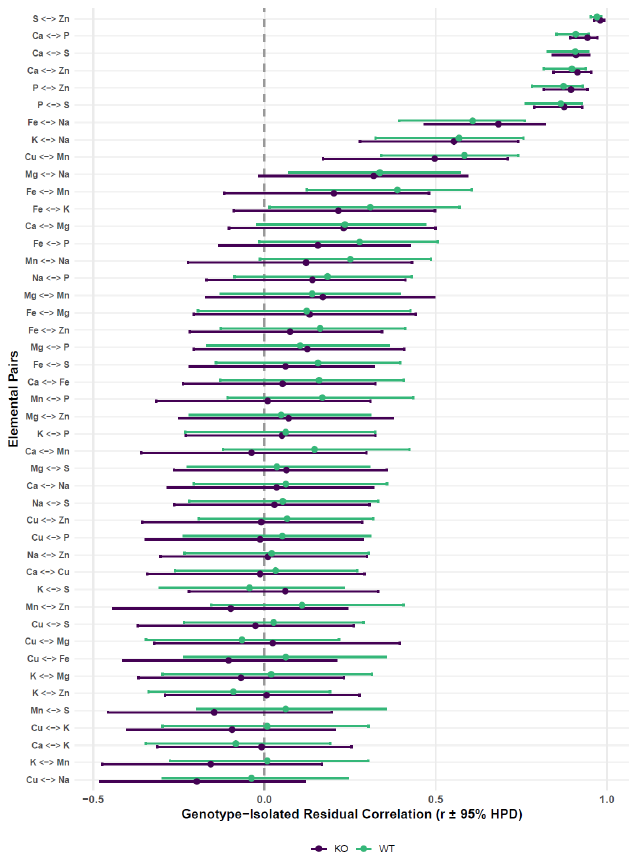

**Figure S5.** Genotype-specific forest plot displaying the posterior median residual correlations and corresponding 95% HPD intervals for all elemental pairs, isolated by genotype. Points represent median and lines represent 95% HPD.

**Table 1**. Global multivariate Bayesian model parameter coordinates and chain diagnostics. Posterior summaries (median estimates, standard errors, and 95% Bayesian Credible Intervals) for all fixed-effect parameters, residual standard deviations (sigma), and residual correlations (rescor) across the 10-element ionomic suite. Chain convergence and sampling efficiency are verified via the Gelman-Rubin diagnostic (Rhat) alongside bulk and tail Effective Sample Sizes (Bulk_ESS, Tail_ESS).

| **Parameter** | **Estimate** | **Std_Error** | **Lower_CI** | **Upper_CI** | **Rhat** | **Bulk_ESS** | **Tail_ESS** |
| --- | --- | --- | --- | --- | --- | --- | --- |
| Ca_Intercept | -3.8195 | 0.7899 | -5.402 | -2.2835 | 1.0033 | 620.0355 | 1336.399 |
| Cu_Intercept | -10.1504 | 0.0709 | -10.2844 | -10.01 | 1.0007 | 1694.6574 | 2330.293 |
| Fe_Intercept | -7.7148 | 0.1551 | -8.0157 | -7.412 | 1.0007 | 1439.2892 | 2605.51 |
| K_Intercept | -4.272 | 0.1558 | -4.574 | -3.9658 | 1.0006 | 1853.0846 | 2392.295 |
| Mg_Intercept | -5.4851 | 0.1044 | -5.6911 | -5.2744 | 0.9998 | 1224.6852 | 2118.526 |
| Mn_Intercept | -8.6652 | 0.0936 | -8.8497 | -8.4762 | 1.0018 | 1420.4715 | 2672.153 |
| Na_Intercept | -4.3052 | 0.1656 | -4.6269 | -3.9804 | 1.0001 | 1230.444 | 2206.672 |
| P_Intercept | -3.8118 | 0.5786 | -4.9543 | -2.6608 | 1.0025 | 658.4672 | 1563.842 |
| S_Intercept | -4.958 | 0.5763 | -6.0881 | -3.8239 | 1.0037 | 604.1112 | 1296.466 |
| Zn_Intercept | -8.2915 | 0.575 | -9.434 | -7.1783 | 1.0037 | 595.1199 | 1249.895 |
| Ca_WT_KOWT | 0.0116 | 1.0561 | -2.0681 | 2.1078 | 1.0054 | 514.108 | 880.9761 |
| Ca_TissueFeces | 0.5692 | 1.1182 | -1.6147 | 2.7903 | 1.0035 | 788.0672 | 1646.718 |
| Ca_TissueFur | -1.9517 | 1.102 | -4.1762 | 0.1272 | 1.0044 | 770.8803 | 1610.067 |
| Ca_TissueOlfactoryBulb | -2.6599 | 1.1087 | -4.8415 | -0.4995 | 1.0027 | 813.6033 | 1560.684 |
| Ca_TissuePMHTH | -5.2653 | 1.1118 | -7.42 | -3.012 | 1.0016 | 795.4002 | 1628.528 |
| Ca_TissueStriatum | -2.5998 | 1.1163 | -4.7434 | -0.4049 | 1.0017 | 888.7187 | 1798.118 |
| Ca_WT_KOWT:TissueFeces | 0.1276 | 1.4794 | -2.8439 | 3.0435 | 1.0026 | 685.0571 | 1117.908 |
| Ca_WT_KOWT:TissueFur | -0.8363 | 1.47 | -3.5854 | 2.0454 | 1.0062 | 712.486 | 1804.084 |
| Ca_WT_KOWT:TissueOlfactoryBulb | -0.9286 | 1.4759 | -3.819 | 1.8934 | 1.0055 | 693.7081 | 1478.354 |
| Ca_WT_KOWT:TissuePMHTH | 1.3164 | 1.4939 | -1.6402 | 4.209 | 1.0017 | 682.7946 | 1437.013 |
| Ca_WT_KOWT:TissueStriatum | -0.0289 | 1.4786 | -2.8531 | 2.8206 | 1.0029 | 770.0914 | 1552.002 |
| Cu_WT_KOWT | -0.0432 | 0.0952 | -0.2317 | 0.1432 | 1.0018 | 1540.2325 | 1904.2 |
| Cu_TissueFeces | 0.2747 | 0.0937 | 0.0888 | 0.4542 | 1.0031 | 1914.9638 | 2417.802 |
| Cu_TissueFur | -1.3302 | 0.095 | -1.5175 | -1.1415 | 1.0009 | 1960.5742 | 2385.81 |
| Cu_TissueOlfactoryBulb | -0.4457 | 0.095 | -0.632 | -0.262 | 1.0009 | 2105.6246 | 2706.541 |
| Cu_TissuePMHTH | -0.7564 | 0.0974 | -0.9442 | -0.5658 | 1.0005 | 2020.4945 | 2441.116 |
| Cu_TissueStriatum | -0.241 | 0.095 | -0.4323 | -0.0562 | 1.0004 | 2185.8326 | 2729.217 |
| Cu_WT_KOWT:TissueFeces | 0.1856 | 0.127 | -0.057 | 0.4348 | 1.0026 | 1830.8307 | 2399.913 |
| Cu_WT_KOWT:TissueFur | 0.0506 | 0.1266 | -0.191 | 0.3009 | 1.0013 | 1855.6192 | 2245.875 |
| Cu_WT_KOWT:TissueOlfactoryBulb | 0.0652 | 0.1271 | -0.1898 | 0.319 | 1.0005 | 1872.4377 | 2485.495 |
| Cu_WT_KOWT:TissuePMHTH | 0.0456 | 0.1287 | -0.2033 | 0.3058 | 1.001 | 1864.625 | 2284.994 |
| Cu_WT_KOWT:TissueStriatum | 0.0387 | 0.127 | -0.2122 | 0.2857 | 1.001 | 1878.4523 | 2138.628 |
| Fe_WT_KOWT | -0.1332 | 0.2056 | -0.5375 | 0.2685 | 1 | 1404.0778 | 2264.51 |
| Fe_TissueFeces | 0.3194 | 0.2142 | -0.1121 | 0.7335 | 1.0011 | 1864.0136 | 2586.191 |
| Fe_TissueFur | -1.9348 | 0.2168 | -2.3506 | -1.4993 | 1.0016 | 1941.6704 | 2709.949 |
| Fe_TissueOlfactoryBulb | -1.0264 | 0.2143 | -1.4438 | -0.5974 | 1.0013 | 1905.7671 | 2639.215 |
| Fe_TissuePMHTH | -1.6111 | 0.2172 | -2.0314 | -1.1826 | 1.0002 | 1986.9563 | 2825.706 |
| Fe_TissueStriatum | -1.4704 | 0.2171 | -1.9039 | -1.0656 | 1.0014 | 1939.2883 | 3006.551 |
| Fe_WT_KOWT:TissueFeces | 0.2414 | 0.2852 | -0.3293 | 0.7832 | 1.0004 | 1854.7226 | 2826.112 |
| Fe_WT_KOWT:TissueFur | 0.1772 | 0.2807 | -0.3784 | 0.7388 | 1.0008 | 1936.4972 | 2975.17 |
| Fe_WT_KOWT:TissueOlfactoryBulb | -0.017 | 0.2854 | -0.5668 | 0.5631 | 1.0007 | 1735.3372 | 2546.263 |
| Fe_WT_KOWT:TissuePMHTH | 0.0445 | 0.285 | -0.5049 | 0.6136 | 0.9997 | 2032.1013 | 2797.804 |
| Fe_WT_KOWT:TissueStriatum | 0.0327 | 0.2854 | -0.5231 | 0.5758 | 1.001 | 1960.1504 | 2824.806 |
| K_WT_KOWT | -0.1012 | 0.2036 | -0.5171 | 0.3008 | 1.0011 | 1755.2832 | 2207.1 |
| K_TissueFeces | -1.1573 | 0.2111 | -1.5712 | -0.753 | 1 | 2322.8689 | 2533.68 |
| K_TissueFur | -1.3476 | 0.2192 | -1.777 | -0.9121 | 1.0003 | 2207.6541 | 2649.155 |
| K_TissueOlfactoryBulb | 0.1693 | 0.2122 | -0.2509 | 0.5815 | 1.0003 | 2260.8521 | 2787.318 |
| K_TissuePMHTH | 0.2637 | 0.2139 | -0.1674 | 0.6794 | 1.0011 | 2335.3839 | 3017.55 |
| K_TissueStriatum | 0.055 | 0.209 | -0.353 | 0.458 | 1.001 | 2298.6282 | 2987.813 |
| K_WT_KOWT:TissueFeces | -0.1765 | 0.2786 | -0.7263 | 0.3851 | 1.0005 | 2276.2874 | 2406.575 |
| K_WT_KOWT:TissueFur | 0.578 | 0.2789 | 0.0241 | 1.1329 | 1.0005 | 2158.065 | 2687.179 |
| K_WT_KOWT:TissueOlfactoryBulb | 0.1837 | 0.2805 | -0.3439 | 0.7326 | 1.0004 | 2254.5602 | 2798.571 |
| K_WT_KOWT:TissuePMHTH | 0.1046 | 0.2784 | -0.4331 | 0.6404 | 1.0006 | 2282.6744 | 2501.399 |
| K_WT_KOWT:TissueStriatum | 0.0466 | 0.2784 | -0.5039 | 0.5855 | 1.0009 | 2142.833 | 2819.82 |
| Mg_WT_KOWT | -0.0674 | 0.1382 | -0.3375 | 0.2079 | 1.0009 | 1155.648 | 1891.328 |
| Mg_TissueFeces | 2.7812 | 0.1464 | 2.4951 | 3.0733 | 1.0007 | 1809.9508 | 2331.074 |
| Mg_TissueFur | -2.4579 | 0.1461 | -2.7476 | -2.1651 | 1.0004 | 1911.005 | 2679.235 |
| Mg_TissueOlfactoryBulb | -1.3863 | 0.1445 | -1.6725 | -1.1098 | 1.0005 | 1734.6746 | 2615.619 |
| Mg_TissuePMHTH | -1.7893 | 0.1468 | -2.0768 | -1.5106 | 0.9995 | 1741.1986 | 2712.655 |
| Mg_TissueStriatum | -0.3086 | 0.1432 | -0.589 | -0.0265 | 1.0009 | 1605.9553 | 2413.993 |
| Mg_WT_KOWT:TissueFeces | 0.3675 | 0.1945 | -0.0112 | 0.7479 | 1.0003 | 1708.9537 | 2316.477 |
| Mg_WT_KOWT:TissueFur | 0.1676 | 0.1909 | -0.2151 | 0.547 | 1.0001 | 1877.2778 | 2731.556 |
| Mg_WT_KOWT:TissueOlfactoryBulb | -0.1031 | 0.1897 | -0.4706 | 0.2596 | 1.0007 | 1594.842 | 2577.253 |
| Mg_WT_KOWT:TissuePMHTH | 0.1625 | 0.1922 | -0.2123 | 0.5436 | 0.9999 | 1432.5786 | 2421.714 |
| Mg_WT_KOWT:TissueStriatum | 0.01 | 0.1946 | -0.3731 | 0.3853 | 1.0001 | 1530.2677 | 2819.808 |
| Mn_WT_KOWT | -0.0272 | 0.1234 | -0.2811 | 0.2139 | 1.0021 | 1217.562 | 2480.42 |
| Mn_TissueFeces | 0.5243 | 0.1271 | 0.271 | 0.7677 | 1.0022 | 1795.6735 | 2570.316 |
| Mn_TissueFur | -4.9426 | 0.128 | -5.2004 | -4.6992 | 1.0014 | 1981.7199 | 2876.066 |
| Mn_TissueOlfactoryBulb | -4.1559 | 0.1274 | -4.3949 | -3.9007 | 1.0014 | 1817.7843 | 2692.256 |
| Mn_TissuePMHTH | -4.5537 | 0.128 | -4.8043 | -4.3043 | 1.0026 | 1965.3949 | 2804.134 |
| Mn_TissueStriatum | -4.3452 | 0.1264 | -4.585 | -4.0977 | 1.0008 | 1878.6277 | 2722.816 |
| Mn_WT_KOWT:TissueFeces | 0.0989 | 0.1688 | -0.2284 | 0.4289 | 1.002 | 1681.0534 | 2182.822 |
| Mn_WT_KOWT:TissueFur | 0.0649 | 0.1692 | -0.2634 | 0.3977 | 1.0018 | 1921.7032 | 2795.561 |
| Mn_WT_KOWT:TissueOlfactoryBulb | -0.0459 | 0.1682 | -0.3715 | 0.2709 | 1.0012 | 1718.0982 | 2405.629 |
| Mn_WT_KOWT:TissuePMHTH | 0.0104 | 0.1666 | -0.3098 | 0.3499 | 1.0016 | 1589.4627 | 2362.028 |
| Mn_WT_KOWT:TissueStriatum | 0.068 | 0.1698 | -0.2682 | 0.4064 | 1.0014 | 1644.8358 | 2184.669 |
| Na_WT_KOWT | -0.0673 | 0.2213 | -0.484 | 0.3675 | 1.0003 | 1146.4071 | 2363.478 |
| Na_TissueFeces | -0.7502 | 0.2333 | -1.2138 | -0.3055 | 1.0003 | 1793.9025 | 2636.974 |
| Na_TissueFur | -2.6984 | 0.2393 | -3.1674 | -2.2259 | 0.9999 | 1733.8638 | 2683.551 |
| Na_TissueOlfactoryBulb | -0.8628 | 0.2327 | -1.3256 | -0.4068 | 1.0006 | 1664.3903 | 2360.782 |
| Na_TissuePMHTH | -0.8205 | 0.2353 | -1.2828 | -0.3678 | 1.001 | 1633.1856 | 2396.464 |
| Na_TissueStriatum | 0.1073 | 0.2302 | -0.3452 | 0.5468 | 1.0001 | 1722.5393 | 2514.148 |
| Na_WT_KOWT:TissueFeces | 0.5394 | 0.3099 | -0.0636 | 1.1518 | 1.0002 | 1709.883 | 2401.132 |
| Na_WT_KOWT:TissueFur | 0.438 | 0.3105 | -0.1801 | 1.0375 | 1.0002 | 1572.8063 | 2321.014 |
| Na_WT_KOWT:TissueOlfactoryBulb | 0.0974 | 0.3087 | -0.5042 | 0.7003 | 1.0007 | 1465.6519 | 2508.161 |
| Na_WT_KOWT:TissuePMHTH | 0.061 | 0.3145 | -0.5431 | 0.6667 | 1.0002 | 1458.4902 | 2432.238 |
| Na_WT_KOWT:TissueStriatum | -0.0128 | 0.3056 | -0.6238 | 0.5763 | 0.9999 | 1584.7522 | 2616.404 |
| P_WT_KOWT | 0.0683 | 0.7773 | -1.436 | 1.6142 | 1.0038 | 535.3787 | 1368.687 |
| P_TissueFeces | 0.211 | 0.8165 | -1.3508 | 1.8179 | 1.0021 | 838.3926 | 1727.86 |
| P_TissueFur | -4.3736 | 0.8093 | -6.0009 | -2.8143 | 1.0034 | 831.2168 | 1456.721 |
| P_TissueOlfactoryBulb | -1.8154 | 0.8178 | -3.4224 | -0.2218 | 1.0015 | 876.4328 | 1712.833 |
| P_TissuePMHTH | -2.8582 | 0.816 | -4.4299 | -1.2247 | 1.0013 | 831.0378 | 1800.38 |
| P_TissueStriatum | -0.1184 | 0.8145 | -1.7394 | 1.5298 | 1.001 | 912.7524 | 1981.541 |
| P_WT_KOWT:TissueFeces | 0.0313 | 1.0885 | -2.1272 | 2.1942 | 1.0019 | 717.2074 | 1476.447 |
| P_WT_KOWT:TissueFur | -0.043 | 1.0716 | -2.0729 | 2.1081 | 1.0051 | 723.2841 | 1528.695 |
| P_WT_KOWT:TissueOlfactoryBulb | -0.1624 | 1.0831 | -2.2731 | 1.9404 | 1.0032 | 726.8771 | 1411.4 |
| P_WT_KOWT:TissuePMHTH | 0.9211 | 1.0986 | -1.2702 | 2.9968 | 1.0016 | 701.4973 | 1681.048 |
| P_WT_KOWT:TissueStriatum | -0.1203 | 1.0838 | -2.2428 | 1.982 | 1.0022 | 775.5888 | 1903.744 |
| S_WT_KOWT | 0.1228 | 0.7833 | -1.4142 | 1.6577 | 1.0048 | 528.9153 | 978.1764 |
| S_TissueFeces | -0.2436 | 0.8116 | -1.8694 | 1.3458 | 1.0023 | 806.7263 | 1576.394 |
| S_TissueFur | 0.4959 | 0.8036 | -1.1071 | 2.0562 | 1.0036 | 723.8297 | 1162.373 |
| S_TissueOlfactoryBulb | -1.3528 | 0.815 | -2.8851 | 0.2781 | 1.0025 | 828.8607 | 1541.881 |
| S_TissuePMHTH | -2.1542 | 0.8135 | -3.7491 | -0.517 | 1.0028 | 783.6307 | 1530.813 |
| S_TissueStriatum | 0.2237 | 0.8083 | -1.3569 | 1.8195 | 1.0015 | 846.3193 | 1771.942 |
| S_WT_KOWT:TissueFeces | -0.132 | 1.0851 | -2.2865 | 1.9649 | 1.0019 | 745.2086 | 1699.604 |
| S_WT_KOWT:TissueFur | -0.471 | 1.0908 | -2.5561 | 1.7132 | 1.0047 | 689.9807 | 1414.147 |
| S_WT_KOWT:TissueOlfactoryBulb | -0.1387 | 1.0842 | -2.2371 | 1.9308 | 1.0036 | 738.9367 | 1516.734 |
| S_WT_KOWT:TissuePMHTH | 0.4554 | 1.1003 | -1.7077 | 2.5732 | 1.0026 | 686.5517 | 1524.568 |
| S_WT_KOWT:TissueStriatum | -0.1671 | 1.0882 | -2.3081 | 1.9521 | 1.0023 | 754.1443 | 1801.679 |
| Zn_WT_KOWT | -0.0458 | 0.7837 | -1.5699 | 1.4929 | 1.0047 | 522.213 | 987.5189 |
| Zn_TissueFeces | 0.1768 | 0.8161 | -1.447 | 1.7507 | 1.0017 | 795.1481 | 1541.763 |
| Zn_TissueFur | -1.3007 | 0.8014 | -2.8779 | 0.2648 | 1.004 | 720.4249 | 1144.445 |
| Zn_TissueOlfactoryBulb | -2.4859 | 0.813 | -4.1233 | -0.8671 | 1.0028 | 803.0538 | 1586.617 |
| Zn_TissuePMHTH | -3.4047 | 0.8164 | -5.0116 | -1.7352 | 1.0025 | 783.9665 | 1657.744 |
| Zn_TissueStriatum | -1.3097 | 0.8055 | -2.8808 | 0.2955 | 1.0017 | 834.537 | 1783.89 |
| Zn_WT_KOWT:TissueFeces | 0.2976 | 1.0844 | -1.8694 | 2.4091 | 1.0023 | 720.7966 | 1569.4 |
| Zn_WT_KOWT:TissueFur | -0.4457 | 1.083 | -2.5459 | 1.7314 | 1.0047 | 682.2006 | 1342.842 |
| Zn_WT_KOWT:TissueOlfactoryBulb | -0.0861 | 1.092 | -2.1961 | 1.9925 | 1.0039 | 727.507 | 1758.748 |
| Zn_WT_KOWT:TissuePMHTH | 0.3449 | 1.1009 | -1.8527 | 2.4613 | 1.0023 | 682.4175 | 1547.812 |
| Zn_WT_KOWT:TissueStriatum | 0.1216 | 1.0869 | -1.9867 | 2.268 | 1.0027 | 745.092 | 1856.264 |
| sigma_Ca | 1.9508 | 0.1404 | 1.6926 | 2.2432 | 1.0005 | 2368.2973 | 2923.016 |
| sigma_Cu | 0.1653 | 0.0152 | 0.1382 | 0.198 | 1 | 4521.2183 | 2928.227 |
| sigma_Fe | 0.3803 | 0.032 | 0.3237 | 0.4492 | 0.9995 | 4939.1726 | 3179.658 |
| sigma_K | 0.3708 | 0.0327 | 0.3132 | 0.4401 | 0.9997 | 4816.255 | 3144.231 |
| sigma_Mg | 0.2475 | 0.022 | 0.2086 | 0.2941 | 1.001 | 4162.1668 | 2996.897 |
| sigma_Mn | 0.2183 | 0.0186 | 0.1854 | 0.2569 | 1.002 | 4548.9127 | 3974.898 |
| sigma_Na | 0.4115 | 0.0342 | 0.3502 | 0.484 | 1.0008 | 4529.0503 | 3505.156 |
| sigma_P | 1.4294 | 0.1061 | 1.24 | 1.6502 | 1.0009 | 2425.3127 | 3324.516 |
| sigma_S | 1.4285 | 0.1053 | 1.2447 | 1.6469 | 1.0023 | 2357.2185 | 2752.953 |
| sigma_Zn | 1.4291 | 0.1053 | 1.2377 | 1.6474 | 1.001 | 2370.7357 | 2856.038 |
| rescor(Ca,Cu) | 0.0028 | 0.0996 | -0.1949 | 0.1963 | 1.0004 | 5712.9917 | 3271.129 |
| rescor(Ca,Fe) | 0.0885 | 0.0968 | -0.1078 | 0.2716 | 0.9999 | 4530.6062 | 3228.501 |
| rescor(Cu,Fe) | -0.0214 | 0.1149 | -0.244 | 0.2077 | 0.9997 | 3845.5973 | 3290.451 |
| rescor(Ca,K) | -0.0512 | 0.0995 | -0.2455 | 0.1457 | 1.0002 | 5208.7795 | 2935.089 |
| rescor(Cu,K) | -0.0347 | 0.1127 | -0.2503 | 0.1871 | 1.0004 | 4599.3376 | 3391.452 |
| rescor(Fe,K) | 0.2344 | 0.1098 | 0.0105 | 0.4356 | 1.0001 | 4553.5183 | 3535.983 |
| rescor(Ca,Mg) | 0.2195 | 0.0969 | 0.0251 | 0.4027 | 1.0007 | 6210.1977 | 3331.619 |
| rescor(Cu,Mg) | -0.0291 | 0.116 | -0.2531 | 0.195 | 1.0008 | 3966.7813 | 3196.274 |
| rescor(Fe,Mg) | 0.1106 | 0.1124 | -0.1111 | 0.3241 | 1.0004 | 4559.4495 | 3795.493 |
| rescor(K,Mg) | -0.028 | 0.1148 | -0.2486 | 0.1964 | 0.9998 | 3987.657 | 3075.311 |
| rescor(Ca,Mn) | 0.0607 | 0.0989 | -0.1433 | 0.2462 | 1.0006 | 5473.4929 | 3446.889 |
| rescor(Cu,Mn) | 0.5192 | 0.089 | 0.3304 | 0.6765 | 1.0001 | 4672.6231 | 3268.483 |
| rescor(Fe,Mn) | 0.2905 | 0.1038 | 0.0802 | 0.4889 | 0.9997 | 3858.0338 | 3414.357 |
| rescor(K,Mn) | -0.0685 | 0.1129 | -0.284 | 0.1518 | 0.9998 | 4546.1182 | 3309.969 |
| rescor(Mg,Mn) | 0.1364 | 0.1134 | -0.0938 | 0.3532 | 1.0002 | 4759.4199 | 3510.005 |
| rescor(Ca,Na) | 0.0434 | 0.0973 | -0.1482 | 0.2298 | 1.0003 | 4282.4536 | 2939.639 |
| rescor(Cu,Na) | -0.1095 | 0.1096 | -0.3221 | 0.109 | 1.0013 | 3619.3563 | 3163.301 |
| rescor(Fe,Na) | 0.6082 | 0.0759 | 0.4453 | 0.7406 | 1.0014 | 4531.2939 | 3423.205 |
| rescor(K,Na) | 0.5319 | 0.0839 | 0.3473 | 0.681 | 0.9999 | 5020.6048 | 3038.682 |
| rescor(Mg,Na) | 0.3088 | 0.1027 | 0.1018 | 0.5023 | 0.9999 | 4389.9981 | 3111.864 |
| rescor(Mn,Na) | 0.1782 | 0.1094 | -0.0393 | 0.3875 | 0.9997 | 4266.4343 | 3314.488 |
| rescor(Ca,P) | 0.9056 | 0.0215 | 0.8579 | 0.9415 | 1.0008 | 2988.3163 | 3328.269 |
| rescor(Cu,P) | 0.0165 | 0.1017 | -0.1806 | 0.2083 | 1.0004 | 5510.8233 | 3826.249 |
| rescor(Fe,P) | 0.2012 | 0.0969 | 4.00E-04 | 0.382 | 0.9995 | 4802.686 | 3579.162 |
| rescor(K,P) | 0.0495 | 0.1022 | -0.1538 | 0.2489 | 1.0007 | 5430.9825 | 3142.553 |
| rescor(Mg,P) | 0.0953 | 0.1014 | -0.1027 | 0.2954 | 1.0013 | 5859.9893 | 3330.294 |
| rescor(Mn,P) | 0.09 | 0.0999 | -0.1102 | 0.2819 | 0.9996 | 5371.5674 | 3177.454 |
| rescor(Na,P) | 0.1547 | 0.0967 | -0.0362 | 0.3399 | 0.9997 | 4471.1311 | 3169.908 |
| rescor(Ca,S) | 0.8842 | 0.0256 | 0.8255 | 0.9254 | 0.9997 | 2795.4782 | 3012.791 |
| rescor(Cu,S) | 0 | 0.0991 | -0.1977 | 0.1912 | 1.0013 | 5244.9627 | 3380.401 |
| rescor(Fe,S) | 0.0935 | 0.0996 | -0.1055 | 0.2793 | 1.0004 | 4434.7831 | 3221.41 |
| rescor(K,S) | 0.0069 | 0.1002 | -0.189 | 0.1974 | 1 | 5201.9324 | 2853.702 |
| rescor(Mg,S) | 0.0296 | 0.1006 | -0.1702 | 0.2227 | 1.0011 | 5943.4632 | 3218.812 |
| rescor(Mn,S) | -0.0321 | 0.1 | -0.2326 | 0.1619 | 1.0021 | 5412.0306 | 3570.957 |
| rescor(Na,S) | 0.032 | 0.0973 | -0.1584 | 0.2178 | 1.0004 | 4655.8477 | 3361.83 |
| rescor(P,S) | 0.8378 | 0.0356 | 0.7559 | 0.8952 | 1.0003 | 2987.3945 | 3392.668 |
| rescor(Ca,Zn) | 0.8801 | 0.0266 | 0.8185 | 0.9229 | 1 | 2644.7602 | 2959.896 |
| rescor(Cu,Zn) | 0.0333 | 0.0998 | -0.1627 | 0.2246 | 1.0005 | 5199.5447 | 3276.531 |
| rescor(Fe,Zn) | 0.1058 | 0.0995 | -0.0952 | 0.2933 | 1.0001 | 4488.6642 | 3445.85 |
| rescor(K,Zn) | -0.0516 | 0.1002 | -0.2456 | 0.1448 | 0.9999 | 5809.9512 | 3446.572 |
| rescor(Mg,Zn) | 0.0437 | 0.1012 | -0.1523 | 0.2388 | 1.0003 | 6074.7492 | 3245.534 |
| rescor(Mn,Zn) | 0.0185 | 0.0997 | -0.1796 | 0.2053 | 1.0009 | 5261.358 | 3645.267 |
| rescor(Na,Zn) | 0.0059 | 0.0978 | -0.1818 | 0.1943 | 0.9997 | 4705.6194 | 3206.292 |
| rescor(P,Zn) | 0.8536 | 0.0322 | 0.7804 | 0.9052 | 1.001 | 3013.7026 | 3234.34 |
| rescor(S,Zn) | 0.9681 | 0.0075 | 0.9518 | 0.9804 | 1.0012 | 2615.2397 | 3225.736 |

**Table 4.** Verification matrix of model predictions. Columns represent predicted median value, lower, and upper quartile, as well as observed median for each tissue, genotype, and element combination.

| **Tissue** | **Element** | **WT_KO** | **Pred_Med** | **Pred_Lwr** | **Pred_Upr** | **Obs_Med** |
| --- | --- | --- | --- | --- | --- | --- |
| Cecal Contents | Ca | WT | -3.80 | -5.15 | -2.45 | -3.79 |
| Cecal Contents | Ca | KO | -3.81 | -5.45 | -2.35 | -3.76 |
| Cecal Contents | Cu | WT | -10.19 | -10.32 | -10.07 | -10.18 |
| Cecal Contents | Cu | KO | -10.15 | -10.29 | -10.01 | -10.19 |
| Cecal Contents | Fe | WT | -7.85 | -8.13 | -7.58 | -7.81 |
| Cecal Contents | Fe | KO | -7.72 | -8.01 | -7.41 | -7.72 |
| Cecal Contents | K | WT | -4.37 | -4.62 | -4.09 | -4.34 |
| Cecal Contents | K | KO | -4.27 | -4.57 | -3.96 | -4.31 |
| Cecal Contents | Mg | WT | -5.55 | -5.72 | -5.38 | -5.48 |
| Cecal Contents | Mg | KO | -5.49 | -5.68 | -5.27 | -5.47 |
| Cecal Contents | Mn | WT | -8.69 | -8.86 | -8.53 | -8.65 |
| Cecal Contents | Mn | KO | -8.67 | -8.85 | -8.48 | -8.69 |
| Cecal Contents | Na | WT | -4.37 | -4.64 | -4.06 | -4.36 |
| Cecal Contents | Na | KO | -4.30 | -4.64 | -3.99 | -4.34 |
| Cecal Contents | P | WT | -3.74 | -4.84 | -2.83 | -3.78 |
| Cecal Contents | P | KO | -3.81 | -4.94 | -2.65 | -3.75 |
| Cecal Contents | S | WT | -4.83 | -5.86 | -3.87 | -4.92 |
| Cecal Contents | S | KO | -4.96 | -6.10 | -3.84 | -4.87 |
| Cecal Contents | Zn | WT | -8.33 | -9.36 | -7.37 | -8.40 |
| Cecal Contents | Zn | KO | -8.31 | -9.40 | -7.16 | -8.26 |
| Feces | Ca | WT | -3.12 | -4.50 | -1.77 | -3.19 |
| Feces | Ca | KO | -3.26 | -4.80 | -1.75 | -3.31 |
| Feces | Cu | WT | -9.73 | -9.85 | -9.62 | -9.60 |
| Feces | Cu | KO | -9.88 | -10.02 | -9.75 | -9.91 |
| Feces | Fe | WT | -7.29 | -7.56 | -7.03 | -7.26 |
| Feces | Fe | KO | -7.40 | -7.68 | -7.08 | -7.37 |
| Feces | K | WT | -5.71 | -5.97 | -5.45 | -5.81 |
| Feces | K | KO | -5.43 | -5.75 | -5.14 | -5.23 |
| Feces | Mg | WT | -2.40 | -2.58 | -2.22 | -2.49 |
| Feces | Mg | KO | -2.70 | -2.92 | -2.52 | -2.71 |
| Feces | Mn | WT | -8.07 | -8.22 | -7.91 | -7.94 |
| Feces | Mn | KO | -8.14 | -8.33 | -7.97 | -8.26 |
| Feces | Na | WT | -4.58 | -4.88 | -4.31 | -4.67 |
| Feces | Na | KO | -5.05 | -5.38 | -4.71 | -4.94 |
| Feces | P | WT | -3.50 | -4.47 | -2.42 | -3.50 |
| Feces | P | KO | -3.60 | -4.74 | -2.46 | -3.57 |
| Feces | S | WT | -5.22 | -6.23 | -4.22 | -5.18 |
| Feces | S | KO | -5.20 | -6.29 | -4.04 | -5.18 |
| Feces | Zn | WT | -7.87 | -8.87 | -6.90 | -7.80 |
| Feces | Zn | KO | -8.13 | -9.22 | -6.98 | -8.16 |
| Fur | Ca | WT | -6.59 | -7.96 | -5.31 | -5.63 |
| Fur | Ca | KO | -5.77 | -7.37 | -4.29 | -5.01 |
| Fur | Cu | WT | -11.47 | -11.59 | -11.35 | -11.46 |
| Fur | Cu | KO | -11.48 | -11.62 | -11.35 | -11.37 |
| Fur | Fe | WT | -9.60 | -9.88 | -9.35 | -9.91 |
| Fur | Fe | KO | -9.65 | -9.94 | -9.34 | -10.07 |
| Fur | K | WT | -5.14 | -5.39 | -4.86 | -4.86 |
| Fur | K | KO | -5.62 | -5.92 | -5.29 | -5.56 |
| Fur | Mg | WT | -7.84 | -8.01 | -7.66 | -7.89 |
| Fur | Mg | KO | -7.94 | -8.15 | -7.75 | -7.96 |
| Fur | Mn | WT | -13.57 | -13.73 | -13.41 | -13.63 |
| Fur | Mn | KO | -13.61 | -13.79 | -13.43 | -13.71 |
| Fur | Na | WT | -6.63 | -6.92 | -6.35 | -6.90 |
| Fur | Na | KO | -7.00 | -7.35 | -6.68 | -7.27 |
| Fur | P | WT | -8.16 | -9.15 | -7.20 | -7.95 |
| Fur | P | KO | -8.18 | -9.31 | -7.01 | -8.08 |
| Fur | S | WT | -4.81 | -5.86 | -3.85 | -3.97 |
| Fur | S | KO | -4.47 | -5.56 | -3.28 | -3.62 |
| Fur | Zn | WT | -10.08 | -11.11 | -9.13 | -9.13 |
| Fur | Zn | KO | -9.59 | -10.70 | -8.43 | -8.78 |
| Olfactory Bulb | Ca | WT | -7.40 | -8.70 | -5.97 | -5.93 |
| Olfactory Bulb | Ca | KO | -6.48 | -8.01 | -4.90 | -4.81 |
| Olfactory Bulb | Cu | WT | -10.57 | -10.69 | -10.45 | -10.57 |
| Olfactory Bulb | Cu | KO | -10.60 | -10.73 | -10.46 | -10.58 |
| Olfactory Bulb | Fe | WT | -8.89 | -9.16 | -8.62 | -8.89 |
| Olfactory Bulb | Fe | KO | -8.74 | -9.05 | -8.44 | -8.76 |
| Olfactory Bulb | K | WT | -4.02 | -4.29 | -3.77 | -4.01 |
| Olfactory Bulb | K | KO | -4.11 | -4.39 | -3.79 | -4.05 |
| Olfactory Bulb | Mg | WT | -7.04 | -7.22 | -6.88 | -7.03 |
| Olfactory Bulb | Mg | KO | -6.87 | -7.07 | -6.68 | -6.96 |
| Olfactory Bulb | Mn | WT | -12.90 | -13.04 | -12.73 | -12.87 |
| Olfactory Bulb | Mn | KO | -12.82 | -13.00 | -12.64 | -12.88 |
| Olfactory Bulb | Na | WT | -5.14 | -5.43 | -4.87 | -5.13 |
| Olfactory Bulb | Na | KO | -5.17 | -5.52 | -4.86 | -5.16 |
| Olfactory Bulb | P | WT | -5.72 | -6.73 | -4.78 | -4.46 |
| Olfactory Bulb | P | KO | -5.62 | -6.82 | -4.54 | -4.10 |
| Olfactory Bulb | S | WT | -6.33 | -7.29 | -5.32 | -5.13 |
| Olfactory Bulb | S | KO | -6.31 | -7.43 | -5.19 | -4.93 |
| Olfactory Bulb | Zn | WT | -10.92 | -11.90 | -9.94 | -9.75 |
| Olfactory Bulb | Zn | KO | -10.77 | -11.94 | -9.71 | -9.45 |
| PMHTH | Ca | WT | -7.75 | -9.16 | -6.47 | -7.93 |
| PMHTH | Ca | KO | -9.10 | -10.62 | -7.57 | -7.57 |
| PMHTH | Cu | WT | -10.90 | -11.03 | -10.79 | -10.90 |
| PMHTH | Cu | KO | -10.91 | -11.06 | -10.78 | -10.94 |
| PMHTH | Fe | WT | -9.42 | -9.68 | -9.16 | -9.47 |
| PMHTH | Fe | KO | -9.32 | -9.65 | -9.03 | -9.32 |
| PMHTH | K | WT | -4.00 | -4.27 | -3.74 | -4.04 |
| PMHTH | K | KO | -4.01 | -4.35 | -3.72 | -3.98 |
| PMHTH | Mg | WT | -7.18 | -7.36 | -7.02 | -7.25 |
| PMHTH | Mg | KO | -7.28 | -7.48 | -7.08 | -7.28 |
| PMHTH | Mn | WT | -13.24 | -13.39 | -13.08 | -13.24 |
| PMHTH | Mn | KO | -13.22 | -13.40 | -13.04 | -13.18 |
| PMHTH | Na | WT | -5.13 | -5.44 | -4.86 | -5.12 |
| PMHTH | Na | KO | -5.13 | -5.47 | -4.80 | -5.11 |
| PMHTH | P | WT | -5.68 | -6.67 | -4.70 | -5.70 |
| PMHTH | P | KO | -6.68 | -7.85 | -5.59 | -5.38 |
| PMHTH | S | WT | -6.53 | -7.58 | -5.61 | -6.43 |
| PMHTH | S | KO | -7.11 | -8.33 | -6.09 | -6.16 |
| PMHTH | Zn | WT | -11.39 | -12.37 | -10.42 | -11.00 |
| PMHTH | Zn | KO | -11.71 | -12.87 | -10.62 | -10.78 |
| Striatum | Ca | WT | -6.43 | -7.71 | -5.10 | -6.52 |
| Striatum | Ca | KO | -6.41 | -7.94 | -4.79 | -6.38 |
| Striatum | Cu | WT | -10.40 | -10.52 | -10.27 | -10.35 |
| Striatum | Cu | KO | -10.39 | -10.53 | -10.26 | -10.41 |
| Striatum | Fe | WT | -9.29 | -9.54 | -9.02 | -9.26 |
| Striatum | Fe | KO | -9.18 | -9.51 | -8.89 | -9.21 |
| Striatum | K | WT | -4.27 | -4.53 | -4.01 | -4.26 |
| Striatum | K | KO | -4.22 | -4.52 | -3.92 | -4.22 |
| Striatum | Mg | WT | -5.85 | -6.03 | -5.67 | -5.88 |
| Striatum | Mg | KO | -5.79 | -6.01 | -5.60 | -5.64 |
| Striatum | Mn | WT | -12.97 | -13.12 | -12.80 | -12.90 |
| Striatum | Mn | KO | -13.01 | -13.19 | -12.83 | -13.02 |
| Striatum | Na | WT | -4.28 | -4.56 | -4.00 | -4.24 |
| Striatum | Na | KO | -4.20 | -4.52 | -3.86 | -4.18 |
| Striatum | P | WT | -3.98 | -4.88 | -2.98 | -3.99 |
| Striatum | P | KO | -3.93 | -5.11 | -2.79 | -3.92 |
| Striatum | S | WT | -4.78 | -5.76 | -3.85 | -4.81 |
| Striatum | S | KO | -4.73 | -5.89 | -3.63 | -4.73 |
| Striatum | Zn | WT | -9.52 | -10.52 | -8.62 | -9.56 |
| Striatum | Zn | KO | -9.60 | -10.71 | -8.48 | -9.66 |

**Table S3.** Post-hoc assessment of multivariate residual dispersion and homoscedasticity. Values represent the Interquartile Range (IQR) and Standard Deviation (SD) of the model-derived residual error on the Additive Log-Ratio (ALR) scale, isolated by genotype.

| **WT_KO** | **Residual_IQR** | **Residual_SD** |
| --- | --- | --- |
| KO | 0.17 | 0.19 |
| WT | 0.21 | 0.23 |

**Table S4.** Post-hoc assessment of multivariate residual dispersion and homoscedasticity. Values represent the Interquartile Range (IQR) and Standard Deviation (SD) of the model-derived residual error on the Additive Log-Ratio (ALR) scale, isolated by tissue type.

| **Tissue** | **Residual_IQR** | **Residual_SD** |
| --- | --- | --- |
| Cecal Contents | 0.155 | 0.157 |
| Feces | 0.197 | 0.240 |
| Fur | 0.154 | 0.145 |
| Olfactory Bulb | 0.119 | 0.113 |
| PMHTH | 0.115 | 0.146 |
| Striatum | 0.528 | 0.392 |

**Table 1**. Global multivariate Bayesian model parameter coordinates and chain diagnostics. Posterior summaries (median estimates, standard errors, and 95% Bayesian Credible Intervals) for all fixed-effect parameters, residual standard deviations (sigma), and residual correlations (rescor) across the 10-element ionomic suite. Chain convergence and sampling efficiency are verified via the Gelman-Rubin diagnostic (Rhat) alongside bulk and tail Effective Sample Sizes (Bulk_ESS, Tail_ESS).

| **Parameter** | **Estimate** | **Std_Error** | **Lower_CI** | **Upper_CI** | **Rhat** | **Bulk_ESS** | **Tail_ESS** |
| --- | --- | --- | --- | --- | --- | --- | --- |
| Ca_Intercept | -3.8195 | 0.7899 | -5.402 | -2.2835 | 1.0033 | 620.0355 | 1336.399 |
| Cu_Intercept | -10.1504 | 0.0709 | -10.2844 | -10.01 | 1.0007 | 1694.6574 | 2330.293 |
| Fe_Intercept | -7.7148 | 0.1551 | -8.0157 | -7.412 | 1.0007 | 1439.2892 | 2605.51 |
| K_Intercept | -4.272 | 0.1558 | -4.574 | -3.9658 | 1.0006 | 1853.0846 | 2392.295 |
| Mg_Intercept | -5.4851 | 0.1044 | -5.6911 | -5.2744 | 0.9998 | 1224.6852 | 2118.526 |
| Mn_Intercept | -8.6652 | 0.0936 | -8.8497 | -8.4762 | 1.0018 | 1420.4715 | 2672.153 |
| Na_Intercept | -4.3052 | 0.1656 | -4.6269 | -3.9804 | 1.0001 | 1230.444 | 2206.672 |
| P_Intercept | -3.8118 | 0.5786 | -4.9543 | -2.6608 | 1.0025 | 658.4672 | 1563.842 |
| S_Intercept | -4.958 | 0.5763 | -6.0881 | -3.8239 | 1.0037 | 604.1112 | 1296.466 |
| Zn_Intercept | -8.2915 | 0.575 | -9.434 | -7.1783 | 1.0037 | 595.1199 | 1249.895 |
| Ca_WT_KOWT | 0.0116 | 1.0561 | -2.0681 | 2.1078 | 1.0054 | 514.108 | 880.9761 |
| Ca_TissueFeces | 0.5692 | 1.1182 | -1.6147 | 2.7903 | 1.0035 | 788.0672 | 1646.718 |
| Ca_TissueFur | -1.9517 | 1.102 | -4.1762 | 0.1272 | 1.0044 | 770.8803 | 1610.067 |
| Ca_TissueOlfactoryBulb | -2.6599 | 1.1087 | -4.8415 | -0.4995 | 1.0027 | 813.6033 | 1560.684 |
| Ca_TissuePMHTH | -5.2653 | 1.1118 | -7.42 | -3.012 | 1.0016 | 795.4002 | 1628.528 |
| Ca_TissueStriatum | -2.5998 | 1.1163 | -4.7434 | -0.4049 | 1.0017 | 888.7187 | 1798.118 |
| Ca_WT_KOWT:TissueFeces | 0.1276 | 1.4794 | -2.8439 | 3.0435 | 1.0026 | 685.0571 | 1117.908 |
| Ca_WT_KOWT:TissueFur | -0.8363 | 1.47 | -3.5854 | 2.0454 | 1.0062 | 712.486 | 1804.084 |
| Ca_WT_KOWT:TissueOlfactoryBulb | -0.9286 | 1.4759 | -3.819 | 1.8934 | 1.0055 | 693.7081 | 1478.354 |
| Ca_WT_KOWT:TissuePMHTH | 1.3164 | 1.4939 | -1.6402 | 4.209 | 1.0017 | 682.7946 | 1437.013 |
| Ca_WT_KOWT:TissueStriatum | -0.0289 | 1.4786 | -2.8531 | 2.8206 | 1.0029 | 770.0914 | 1552.002 |
| Cu_WT_KOWT | -0.0432 | 0.0952 | -0.2317 | 0.1432 | 1.0018 | 1540.2325 | 1904.2 |
| Cu_TissueFeces | 0.2747 | 0.0937 | 0.0888 | 0.4542 | 1.0031 | 1914.9638 | 2417.802 |
| Cu_TissueFur | -1.3302 | 0.095 | -1.5175 | -1.1415 | 1.0009 | 1960.5742 | 2385.81 |
| Cu_TissueOlfactoryBulb | -0.4457 | 0.095 | -0.632 | -0.262 | 1.0009 | 2105.6246 | 2706.541 |
| Cu_TissuePMHTH | -0.7564 | 0.0974 | -0.9442 | -0.5658 | 1.0005 | 2020.4945 | 2441.116 |
| Cu_TissueStriatum | -0.241 | 0.095 | -0.4323 | -0.0562 | 1.0004 | 2185.8326 | 2729.217 |
| Cu_WT_KOWT:TissueFeces | 0.1856 | 0.127 | -0.057 | 0.4348 | 1.0026 | 1830.8307 | 2399.913 |
| Cu_WT_KOWT:TissueFur | 0.0506 | 0.1266 | -0.191 | 0.3009 | 1.0013 | 1855.6192 | 2245.875 |
| Cu_WT_KOWT:TissueOlfactoryBulb | 0.0652 | 0.1271 | -0.1898 | 0.319 | 1.0005 | 1872.4377 | 2485.495 |
| Cu_WT_KOWT:TissuePMHTH | 0.0456 | 0.1287 | -0.2033 | 0.3058 | 1.001 | 1864.625 | 2284.994 |
| Cu_WT_KOWT:TissueStriatum | 0.0387 | 0.127 | -0.2122 | 0.2857 | 1.001 | 1878.4523 | 2138.628 |
| Fe_WT_KOWT | -0.1332 | 0.2056 | -0.5375 | 0.2685 | 1 | 1404.0778 | 2264.51 |
| Fe_TissueFeces | 0.3194 | 0.2142 | -0.1121 | 0.7335 | 1.0011 | 1864.0136 | 2586.191 |
| Fe_TissueFur | -1.9348 | 0.2168 | -2.3506 | -1.4993 | 1.0016 | 1941.6704 | 2709.949 |
| Fe_TissueOlfactoryBulb | -1.0264 | 0.2143 | -1.4438 | -0.5974 | 1.0013 | 1905.7671 | 2639.215 |
| Fe_TissuePMHTH | -1.6111 | 0.2172 | -2.0314 | -1.1826 | 1.0002 | 1986.9563 | 2825.706 |
| Fe_TissueStriatum | -1.4704 | 0.2171 | -1.9039 | -1.0656 | 1.0014 | 1939.2883 | 3006.551 |
| Fe_WT_KOWT:TissueFeces | 0.2414 | 0.2852 | -0.3293 | 0.7832 | 1.0004 | 1854.7226 | 2826.112 |
| Fe_WT_KOWT:TissueFur | 0.1772 | 0.2807 | -0.3784 | 0.7388 | 1.0008 | 1936.4972 | 2975.17 |
| Fe_WT_KOWT:TissueOlfactoryBulb | -0.017 | 0.2854 | -0.5668 | 0.5631 | 1.0007 | 1735.3372 | 2546.263 |
| Fe_WT_KOWT:TissuePMHTH | 0.0445 | 0.285 | -0.5049 | 0.6136 | 0.9997 | 2032.1013 | 2797.804 |
| Fe_WT_KOWT:TissueStriatum | 0.0327 | 0.2854 | -0.5231 | 0.5758 | 1.001 | 1960.1504 | 2824.806 |
| K_WT_KOWT | -0.1012 | 0.2036 | -0.5171 | 0.3008 | 1.0011 | 1755.2832 | 2207.1 |
| K_TissueFeces | -1.1573 | 0.2111 | -1.5712 | -0.753 | 1 | 2322.8689 | 2533.68 |
| K_TissueFur | -1.3476 | 0.2192 | -1.777 | -0.9121 | 1.0003 | 2207.6541 | 2649.155 |
| K_TissueOlfactoryBulb | 0.1693 | 0.2122 | -0.2509 | 0.5815 | 1.0003 | 2260.8521 | 2787.318 |
| K_TissuePMHTH | 0.2637 | 0.2139 | -0.1674 | 0.6794 | 1.0011 | 2335.3839 | 3017.55 |
| K_TissueStriatum | 0.055 | 0.209 | -0.353 | 0.458 | 1.001 | 2298.6282 | 2987.813 |
| K_WT_KOWT:TissueFeces | -0.1765 | 0.2786 | -0.7263 | 0.3851 | 1.0005 | 2276.2874 | 2406.575 |
| K_WT_KOWT:TissueFur | 0.578 | 0.2789 | 0.0241 | 1.1329 | 1.0005 | 2158.065 | 2687.179 |
| K_WT_KOWT:TissueOlfactoryBulb | 0.1837 | 0.2805 | -0.3439 | 0.7326 | 1.0004 | 2254.5602 | 2798.571 |
| K_WT_KOWT:TissuePMHTH | 0.1046 | 0.2784 | -0.4331 | 0.6404 | 1.0006 | 2282.6744 | 2501.399 |
| K_WT_KOWT:TissueStriatum | 0.0466 | 0.2784 | -0.5039 | 0.5855 | 1.0009 | 2142.833 | 2819.82 |
| Mg_WT_KOWT | -0.0674 | 0.1382 | -0.3375 | 0.2079 | 1.0009 | 1155.648 | 1891.328 |
| Mg_TissueFeces | 2.7812 | 0.1464 | 2.4951 | 3.0733 | 1.0007 | 1809.9508 | 2331.074 |
| Mg_TissueFur | -2.4579 | 0.1461 | -2.7476 | -2.1651 | 1.0004 | 1911.005 | 2679.235 |
| Mg_TissueOlfactoryBulb | -1.3863 | 0.1445 | -1.6725 | -1.1098 | 1.0005 | 1734.6746 | 2615.619 |
| Mg_TissuePMHTH | -1.7893 | 0.1468 | -2.0768 | -1.5106 | 0.9995 | 1741.1986 | 2712.655 |
| Mg_TissueStriatum | -0.3086 | 0.1432 | -0.589 | -0.0265 | 1.0009 | 1605.9553 | 2413.993 |
| Mg_WT_KOWT:TissueFeces | 0.3675 | 0.1945 | -0.0112 | 0.7479 | 1.0003 | 1708.9537 | 2316.477 |
| Mg_WT_KOWT:TissueFur | 0.1676 | 0.1909 | -0.2151 | 0.547 | 1.0001 | 1877.2778 | 2731.556 |
| Mg_WT_KOWT:TissueOlfactoryBulb | -0.1031 | 0.1897 | -0.4706 | 0.2596 | 1.0007 | 1594.842 | 2577.253 |
| Mg_WT_KOWT:TissuePMHTH | 0.1625 | 0.1922 | -0.2123 | 0.5436 | 0.9999 | 1432.5786 | 2421.714 |
| Mg_WT_KOWT:TissueStriatum | 0.01 | 0.1946 | -0.3731 | 0.3853 | 1.0001 | 1530.2677 | 2819.808 |
| Mn_WT_KOWT | -0.0272 | 0.1234 | -0.2811 | 0.2139 | 1.0021 | 1217.562 | 2480.42 |
| Mn_TissueFeces | 0.5243 | 0.1271 | 0.271 | 0.7677 | 1.0022 | 1795.6735 | 2570.316 |
| Mn_TissueFur | -4.9426 | 0.128 | -5.2004 | -4.6992 | 1.0014 | 1981.7199 | 2876.066 |
| Mn_TissueOlfactoryBulb | -4.1559 | 0.1274 | -4.3949 | -3.9007 | 1.0014 | 1817.7843 | 2692.256 |
| Mn_TissuePMHTH | -4.5537 | 0.128 | -4.8043 | -4.3043 | 1.0026 | 1965.3949 | 2804.134 |
| Mn_TissueStriatum | -4.3452 | 0.1264 | -4.585 | -4.0977 | 1.0008 | 1878.6277 | 2722.816 |
| Mn_WT_KOWT:TissueFeces | 0.0989 | 0.1688 | -0.2284 | 0.4289 | 1.002 | 1681.0534 | 2182.822 |
| Mn_WT_KOWT:TissueFur | 0.0649 | 0.1692 | -0.2634 | 0.3977 | 1.0018 | 1921.7032 | 2795.561 |
| Mn_WT_KOWT:TissueOlfactoryBulb | -0.0459 | 0.1682 | -0.3715 | 0.2709 | 1.0012 | 1718.0982 | 2405.629 |
| Mn_WT_KOWT:TissuePMHTH | 0.0104 | 0.1666 | -0.3098 | 0.3499 | 1.0016 | 1589.4627 | 2362.028 |
| Mn_WT_KOWT:TissueStriatum | 0.068 | 0.1698 | -0.2682 | 0.4064 | 1.0014 | 1644.8358 | 2184.669 |
| Na_WT_KOWT | -0.0673 | 0.2213 | -0.484 | 0.3675 | 1.0003 | 1146.4071 | 2363.478 |
| Na_TissueFeces | -0.7502 | 0.2333 | -1.2138 | -0.3055 | 1.0003 | 1793.9025 | 2636.974 |
| Na_TissueFur | -2.6984 | 0.2393 | -3.1674 | -2.2259 | 0.9999 | 1733.8638 | 2683.551 |
| Na_TissueOlfactoryBulb | -0.8628 | 0.2327 | -1.3256 | -0.4068 | 1.0006 | 1664.3903 | 2360.782 |
| Na_TissuePMHTH | -0.8205 | 0.2353 | -1.2828 | -0.3678 | 1.001 | 1633.1856 | 2396.464 |
| Na_TissueStriatum | 0.1073 | 0.2302 | -0.3452 | 0.5468 | 1.0001 | 1722.5393 | 2514.148 |
| Na_WT_KOWT:TissueFeces | 0.5394 | 0.3099 | -0.0636 | 1.1518 | 1.0002 | 1709.883 | 2401.132 |
| Na_WT_KOWT:TissueFur | 0.438 | 0.3105 | -0.1801 | 1.0375 | 1.0002 | 1572.8063 | 2321.014 |
| Na_WT_KOWT:TissueOlfactoryBulb | 0.0974 | 0.3087 | -0.5042 | 0.7003 | 1.0007 | 1465.6519 | 2508.161 |
| Na_WT_KOWT:TissuePMHTH | 0.061 | 0.3145 | -0.5431 | 0.6667 | 1.0002 | 1458.4902 | 2432.238 |
| Na_WT_KOWT:TissueStriatum | -0.0128 | 0.3056 | -0.6238 | 0.5763 | 0.9999 | 1584.7522 | 2616.404 |
| P_WT_KOWT | 0.0683 | 0.7773 | -1.436 | 1.6142 | 1.0038 | 535.3787 | 1368.687 |
| P_TissueFeces | 0.211 | 0.8165 | -1.3508 | 1.8179 | 1.0021 | 838.3926 | 1727.86 |
| P_TissueFur | -4.3736 | 0.8093 | -6.0009 | -2.8143 | 1.0034 | 831.2168 | 1456.721 |
| P_TissueOlfactoryBulb | -1.8154 | 0.8178 | -3.4224 | -0.2218 | 1.0015 | 876.4328 | 1712.833 |
| P_TissuePMHTH | -2.8582 | 0.816 | -4.4299 | -1.2247 | 1.0013 | 831.0378 | 1800.38 |
| P_TissueStriatum | -0.1184 | 0.8145 | -1.7394 | 1.5298 | 1.001 | 912.7524 | 1981.541 |
| P_WT_KOWT:TissueFeces | 0.0313 | 1.0885 | -2.1272 | 2.1942 | 1.0019 | 717.2074 | 1476.447 |
| P_WT_KOWT:TissueFur | -0.043 | 1.0716 | -2.0729 | 2.1081 | 1.0051 | 723.2841 | 1528.695 |
| P_WT_KOWT:TissueOlfactoryBulb | -0.1624 | 1.0831 | -2.2731 | 1.9404 | 1.0032 | 726.8771 | 1411.4 |
| P_WT_KOWT:TissuePMHTH | 0.9211 | 1.0986 | -1.2702 | 2.9968 | 1.0016 | 701.4973 | 1681.048 |
| P_WT_KOWT:TissueStriatum | -0.1203 | 1.0838 | -2.2428 | 1.982 | 1.0022 | 775.5888 | 1903.744 |
| S_WT_KOWT | 0.1228 | 0.7833 | -1.4142 | 1.6577 | 1.0048 | 528.9153 | 978.1764 |
| S_TissueFeces | -0.2436 | 0.8116 | -1.8694 | 1.3458 | 1.0023 | 806.7263 | 1576.394 |
| S_TissueFur | 0.4959 | 0.8036 | -1.1071 | 2.0562 | 1.0036 | 723.8297 | 1162.373 |
| S_TissueOlfactoryBulb | -1.3528 | 0.815 | -2.8851 | 0.2781 | 1.0025 | 828.8607 | 1541.881 |
| S_TissuePMHTH | -2.1542 | 0.8135 | -3.7491 | -0.517 | 1.0028 | 783.6307 | 1530.813 |
| S_TissueStriatum | 0.2237 | 0.8083 | -1.3569 | 1.8195 | 1.0015 | 846.3193 | 1771.942 |
| S_WT_KOWT:TissueFeces | -0.132 | 1.0851 | -2.2865 | 1.9649 | 1.0019 | 745.2086 | 1699.604 |
| S_WT_KOWT:TissueFur | -0.471 | 1.0908 | -2.5561 | 1.7132 | 1.0047 | 689.9807 | 1414.147 |
| S_WT_KOWT:TissueOlfactoryBulb | -0.1387 | 1.0842 | -2.2371 | 1.9308 | 1.0036 | 738.9367 | 1516.734 |
| S_WT_KOWT:TissuePMHTH | 0.4554 | 1.1003 | -1.7077 | 2.5732 | 1.0026 | 686.5517 | 1524.568 |
| S_WT_KOWT:TissueStriatum | -0.1671 | 1.0882 | -2.3081 | 1.9521 | 1.0023 | 754.1443 | 1801.679 |
| Zn_WT_KOWT | -0.0458 | 0.7837 | -1.5699 | 1.4929 | 1.0047 | 522.213 | 987.5189 |
| Zn_TissueFeces | 0.1768 | 0.8161 | -1.447 | 1.7507 | 1.0017 | 795.1481 | 1541.763 |
| Zn_TissueFur | -1.3007 | 0.8014 | -2.8779 | 0.2648 | 1.004 | 720.4249 | 1144.445 |
| Zn_TissueOlfactoryBulb | -2.4859 | 0.813 | -4.1233 | -0.8671 | 1.0028 | 803.0538 | 1586.617 |
| Zn_TissuePMHTH | -3.4047 | 0.8164 | -5.0116 | -1.7352 | 1.0025 | 783.9665 | 1657.744 |
| Zn_TissueStriatum | -1.3097 | 0.8055 | -2.8808 | 0.2955 | 1.0017 | 834.537 | 1783.89 |
| Zn_WT_KOWT:TissueFeces | 0.2976 | 1.0844 | -1.8694 | 2.4091 | 1.0023 | 720.7966 | 1569.4 |
| Zn_WT_KOWT:TissueFur | -0.4457 | 1.083 | -2.5459 | 1.7314 | 1.0047 | 682.2006 | 1342.842 |
| Zn_WT_KOWT:TissueOlfactoryBulb | -0.0861 | 1.092 | -2.1961 | 1.9925 | 1.0039 | 727.507 | 1758.748 |
| Zn_WT_KOWT:TissuePMHTH | 0.3449 | 1.1009 | -1.8527 | 2.4613 | 1.0023 | 682.4175 | 1547.812 |
| Zn_WT_KOWT:TissueStriatum | 0.1216 | 1.0869 | -1.9867 | 2.268 | 1.0027 | 745.092 | 1856.264 |
| sigma_Ca | 1.9508 | 0.1404 | 1.6926 | 2.2432 | 1.0005 | 2368.2973 | 2923.016 |
| sigma_Cu | 0.1653 | 0.0152 | 0.1382 | 0.198 | 1 | 4521.2183 | 2928.227 |
| sigma_Fe | 0.3803 | 0.032 | 0.3237 | 0.4492 | 0.9995 | 4939.1726 | 3179.658 |
| sigma_K | 0.3708 | 0.0327 | 0.3132 | 0.4401 | 0.9997 | 4816.255 | 3144.231 |
| sigma_Mg | 0.2475 | 0.022 | 0.2086 | 0.2941 | 1.001 | 4162.1668 | 2996.897 |
| sigma_Mn | 0.2183 | 0.0186 | 0.1854 | 0.2569 | 1.002 | 4548.9127 | 3974.898 |
| sigma_Na | 0.4115 | 0.0342 | 0.3502 | 0.484 | 1.0008 | 4529.0503 | 3505.156 |
| sigma_P | 1.4294 | 0.1061 | 1.24 | 1.6502 | 1.0009 | 2425.3127 | 3324.516 |
| sigma_S | 1.4285 | 0.1053 | 1.2447 | 1.6469 | 1.0023 | 2357.2185 | 2752.953 |
| sigma_Zn | 1.4291 | 0.1053 | 1.2377 | 1.6474 | 1.001 | 2370.7357 | 2856.038 |
| rescor(Ca,Cu) | 0.0028 | 0.0996 | -0.1949 | 0.1963 | 1.0004 | 5712.9917 | 3271.129 |
| rescor(Ca,Fe) | 0.0885 | 0.0968 | -0.1078 | 0.2716 | 0.9999 | 4530.6062 | 3228.501 |
| rescor(Cu,Fe) | -0.0214 | 0.1149 | -0.244 | 0.2077 | 0.9997 | 3845.5973 | 3290.451 |
| rescor(Ca,K) | -0.0512 | 0.0995 | -0.2455 | 0.1457 | 1.0002 | 5208.7795 | 2935.089 |
| rescor(Cu,K) | -0.0347 | 0.1127 | -0.2503 | 0.1871 | 1.0004 | 4599.3376 | 3391.452 |
| rescor(Fe,K) | 0.2344 | 0.1098 | 0.0105 | 0.4356 | 1.0001 | 4553.5183 | 3535.983 |
| rescor(Ca,Mg) | 0.2195 | 0.0969 | 0.0251 | 0.4027 | 1.0007 | 6210.1977 | 3331.619 |
| rescor(Cu,Mg) | -0.0291 | 0.116 | -0.2531 | 0.195 | 1.0008 | 3966.7813 | 3196.274 |
| rescor(Fe,Mg) | 0.1106 | 0.1124 | -0.1111 | 0.3241 | 1.0004 | 4559.4495 | 3795.493 |
| rescor(K,Mg) | -0.028 | 0.1148 | -0.2486 | 0.1964 | 0.9998 | 3987.657 | 3075.311 |
| rescor(Ca,Mn) | 0.0607 | 0.0989 | -0.1433 | 0.2462 | 1.0006 | 5473.4929 | 3446.889 |
| rescor(Cu,Mn) | 0.5192 | 0.089 | 0.3304 | 0.6765 | 1.0001 | 4672.6231 | 3268.483 |
| rescor(Fe,Mn) | 0.2905 | 0.1038 | 0.0802 | 0.4889 | 0.9997 | 3858.0338 | 3414.357 |
| rescor(K,Mn) | -0.0685 | 0.1129 | -0.284 | 0.1518 | 0.9998 | 4546.1182 | 3309.969 |
| rescor(Mg,Mn) | 0.1364 | 0.1134 | -0.0938 | 0.3532 | 1.0002 | 4759.4199 | 3510.005 |
| rescor(Ca,Na) | 0.0434 | 0.0973 | -0.1482 | 0.2298 | 1.0003 | 4282.4536 | 2939.639 |
| rescor(Cu,Na) | -0.1095 | 0.1096 | -0.3221 | 0.109 | 1.0013 | 3619.3563 | 3163.301 |
| rescor(Fe,Na) | 0.6082 | 0.0759 | 0.4453 | 0.7406 | 1.0014 | 4531.2939 | 3423.205 |
| rescor(K,Na) | 0.5319 | 0.0839 | 0.3473 | 0.681 | 0.9999 | 5020.6048 | 3038.682 |
| rescor(Mg,Na) | 0.3088 | 0.1027 | 0.1018 | 0.5023 | 0.9999 | 4389.9981 | 3111.864 |
| rescor(Mn,Na) | 0.1782 | 0.1094 | -0.0393 | 0.3875 | 0.9997 | 4266.4343 | 3314.488 |
| rescor(Ca,P) | 0.9056 | 0.0215 | 0.8579 | 0.9415 | 1.0008 | 2988.3163 | 3328.269 |
| rescor(Cu,P) | 0.0165 | 0.1017 | -0.1806 | 0.2083 | 1.0004 | 5510.8233 | 3826.249 |
| rescor(Fe,P) | 0.2012 | 0.0969 | 4.00E-04 | 0.382 | 0.9995 | 4802.686 | 3579.162 |
| rescor(K,P) | 0.0495 | 0.1022 | -0.1538 | 0.2489 | 1.0007 | 5430.9825 | 3142.553 |
| rescor(Mg,P) | 0.0953 | 0.1014 | -0.1027 | 0.2954 | 1.0013 | 5859.9893 | 3330.294 |
| rescor(Mn,P) | 0.09 | 0.0999 | -0.1102 | 0.2819 | 0.9996 | 5371.5674 | 3177.454 |
| rescor(Na,P) | 0.1547 | 0.0967 | -0.0362 | 0.3399 | 0.9997 | 4471.1311 | 3169.908 |
| rescor(Ca,S) | 0.8842 | 0.0256 | 0.8255 | 0.9254 | 0.9997 | 2795.4782 | 3012.791 |
| rescor(Cu,S) | 0 | 0.0991 | -0.1977 | 0.1912 | 1.0013 | 5244.9627 | 3380.401 |
| rescor(Fe,S) | 0.0935 | 0.0996 | -0.1055 | 0.2793 | 1.0004 | 4434.7831 | 3221.41 |
| rescor(K,S) | 0.0069 | 0.1002 | -0.189 | 0.1974 | 1 | 5201.9324 | 2853.702 |
| rescor(Mg,S) | 0.0296 | 0.1006 | -0.1702 | 0.2227 | 1.0011 | 5943.4632 | 3218.812 |
| rescor(Mn,S) | -0.0321 | 0.1 | -0.2326 | 0.1619 | 1.0021 | 5412.0306 | 3570.957 |
| rescor(Na,S) | 0.032 | 0.0973 | -0.1584 | 0.2178 | 1.0004 | 4655.8477 | 3361.83 |
| rescor(P,S) | 0.8378 | 0.0356 | 0.7559 | 0.8952 | 1.0003 | 2987.3945 | 3392.668 |
| rescor(Ca,Zn) | 0.8801 | 0.0266 | 0.8185 | 0.9229 | 1 | 2644.7602 | 2959.896 |
| rescor(Cu,Zn) | 0.0333 | 0.0998 | -0.1627 | 0.2246 | 1.0005 | 5199.5447 | 3276.531 |
| rescor(Fe,Zn) | 0.1058 | 0.0995 | -0.0952 | 0.2933 | 1.0001 | 4488.6642 | 3445.85 |
| rescor(K,Zn) | -0.0516 | 0.1002 | -0.2456 | 0.1448 | 0.9999 | 5809.9512 | 3446.572 |
| rescor(Mg,Zn) | 0.0437 | 0.1012 | -0.1523 | 0.2388 | 1.0003 | 6074.7492 | 3245.534 |
| rescor(Mn,Zn) | 0.0185 | 0.0997 | -0.1796 | 0.2053 | 1.0009 | 5261.358 | 3645.267 |
| rescor(Na,Zn) | 0.0059 | 0.0978 | -0.1818 | 0.1943 | 0.9997 | 4705.6194 | 3206.292 |
| rescor(P,Zn) | 0.8536 | 0.0322 | 0.7804 | 0.9052 | 1.001 | 3013.7026 | 3234.34 |
| rescor(S,Zn) | 0.9681 | 0.0075 | 0.9518 | 0.9804 | 1.0012 | 2615.2397 | 3225.736 |
